# Predictive link between systemic metabolism and cytokine signatures in the brain of apolipoprotein E ε4 mice

**DOI:** 10.1101/2022.02.11.480074

**Authors:** Rebecca M Fleeman, Amanda M Snyder, Madison K Kuhn, Dennis C Chan, Grace C Smith, Nicole A Crowley, Amy C Arnold, Elizabeth A Proctor

## Abstract

The ε4 variant of apolipoprotein E (*APOE*) is the strongest and most common genetic risk factor for Alzheimer’s disease (AD). While the mechanism of conveyed risk is incompletely understood, promotion of inflammation, dysregulated metabolism, and protein misfolding and aggregation are contributors to accelerating disease. Here we determined the concurrent effects of systemic metabolic changes and brain inflammation in young (3-month-old) and aged (18-month-old) male and female mice carrying the *APOE4* gene. Using functional metabolic assays alongside multivariate modeling of hippocampal cytokine levels, we found that brain cytokine signatures are predictive of systemic metabolic outcomes, independent of AD proteinopathies. Male and female mice each produce different cytokine signatures as they age and as their systemic metabolic phenotype declines, and these signatures are *APOE* genotype dependent. Ours is the first study to identify a quantitative and predictive link between systemic metabolism and specific pathological cytokine signatures in the brain. Our results highlight the effects of APOE4 beyond the brain and suggest the potential for bi-directional influence of risk factors in the brain and periphery.

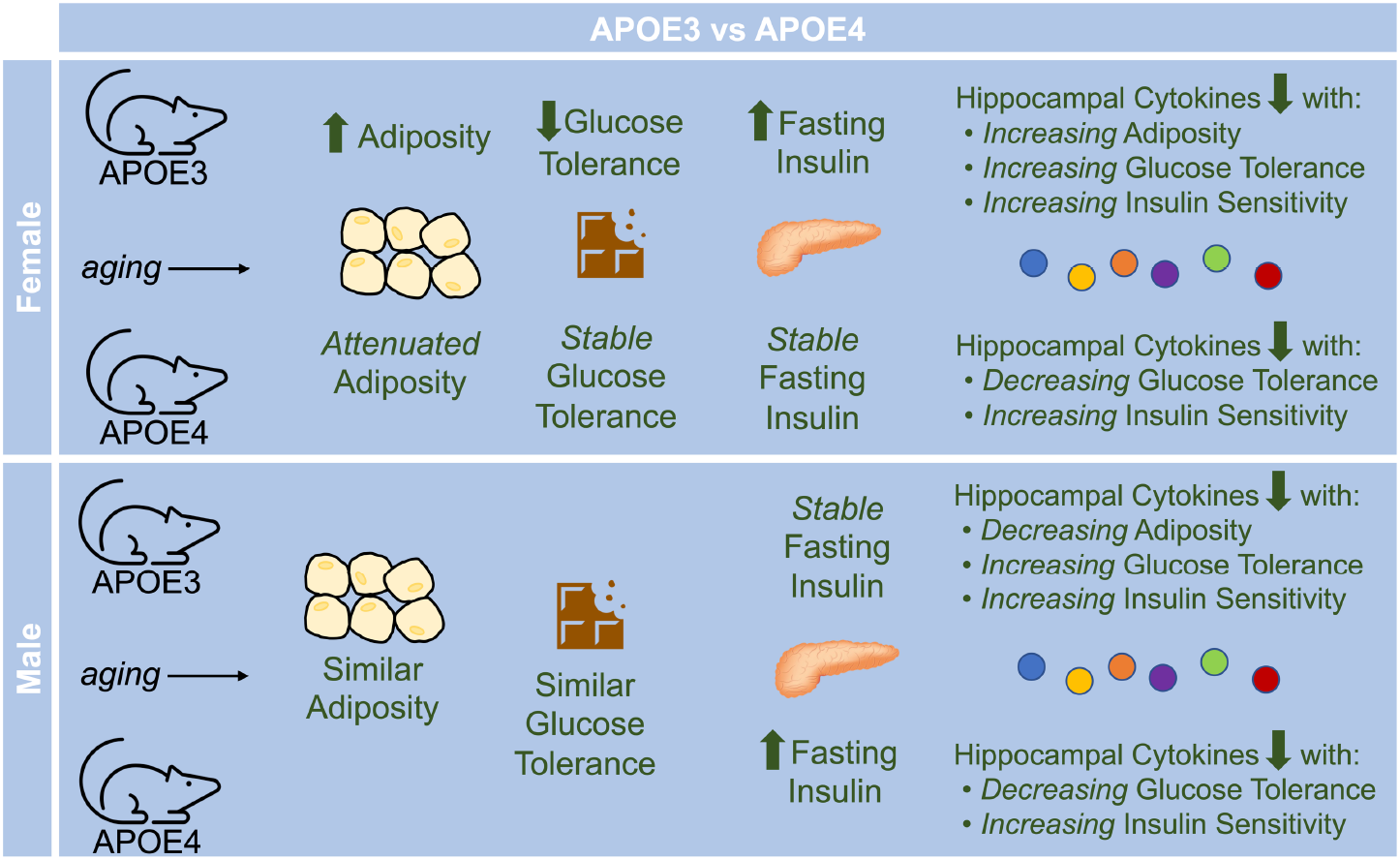

## 1. Introduction

Over 6.2 million Americans are currently living with Alzheimer’s disease (AD), a neurodegenerative disease manifesting as cognitive decline and memory loss(Alzheimer’s Association, 2021). While aging is the greatest overall risk factor for development of AD, the strongest and most common genetic risk factor is the ε4 variant of apolipoprotein E (APOE4)(Corder et al., 1993). The *APOE* gene has three isoforms, ε2, ε3, and ε4, with allelic frequencies of 7%, 79%, and 14% in the US population, respectively(Lanfranco et al., 2020). *APOE4* carriers have a dose-dependent 3-14-fold increased risk over *APOE3* carriers for developing AD(Lanfranco et al., 2020); however, the physiological mechanism of this conveyed risk is not yet fully understood.

In addition to promoting onset and progression of AD, *APOE4* is associated with inflammatory processes and outcomes in both the brain and the periphery, including increased low-density lipoprotein (LDL) cholesterol, increased incidence of cardiovascular disease, and promotion of metabolic syndrome(Bennet et al., 2007; El-Lebedy et al., 2016; Lagging et al., 2019; Lahoz et al., 2001; Torres-Perez et al., 2016; Tsuang et al., 2013). As a genetic variant, *APOE4* is a risk factor that carriers possess from birth, with effects potentially building over their lifetime(Fernandez et al., 2019). Thus, many *APOE4*-related changes, such as increased inflammation(Jofre-Monseny et al., 2008; Wang et al., 2020) and metabolic dysfunction(Jagust et al., 2012; Wu et al., 2018; Zhao et al., 2017), likely occur before the age at which the hallmark AD proteinopathies, amyloid-β (Aβ) plaques and neurofibrillary tau tangles, form in the brain. However, the majority of the focus for APOE AD research has been on the interaction of APOE4 with Aβ and tau, despite the appearance of appreciable amounts of these pathological proteins only later in life(Huynh et al., 2017; Litvinchuk et al., 2021; Liu et al., 2017; Martínez-Martínez et al., 2020; Shi et al., 2017). We propose that, because APOE4 is present from birth and has been shown to have deleterious effects on immune function and metabolism well before the age of AD onset(Flowers and Rebeck, 2020), long-term APOE4-driven systemic and brain immunometabolic effects can provoke an AD-inducible environment independent of interactions with Aβ or tau.

APOE4 alters lipid(Lin et al., 2018; Vardarajan et al., 2020), cholesterol(Lin et al., 2018), and glucose metabolism in the brain(Farmer et al., 2021; Qi et al., 2021; Wu et al., 2018; Zhao et al., 2017), yet little is known about the influence of systemic metabolic function on molecular signaling networks in the brain in the context of APOE4(Christensen and Pike, 2019; Jones et al., 2019; Moser and Pike, 2017). Previous studies have identified a link between systemic metabolism and hippocampal electrophysiology changes(Seto et al., 1983; Tingley et al., 2021), where communication between the hippocampus and periphery can involve hippocampal signaling to the pancreas and liver, as well as to the hypothalamus(Lathe, 2001; Tingley et al., 2021), a major metabolic control center in the brain. A recent study from Guojun Bu’s group even identified that liver-expressed APOE in mice could affect cognition and synaptic function(Golden and Johnson, 2022; Liu et al., 2022), indicating the importance of peripheral APOE effects on the brain. An approach that identifies the relationships of APOE4 effects on systemic metabolic function and neuroinflammation together, instead of studying the brain and periphery as isolated systems, is necessary to define the disease-promoting environment created by the interacting effects of systemic metabolic dysfunction and neuroinflammation under the combined influence of *APOE* genotype and aging.

Here, we determine that aging-related cytokine patterns in the hippocampus can predict systemic metabolic outcomes of young and aged humanized APOE3 and APOE4 male and female mice. Specifically, we uncover unique patterns of cytokines in APOE3 versus APOE4 mice that correlate with body adiposity, glucose tolerance, and insulin sensitivity. Male and female mice exhibit differing cytokine signatures that correlate with peripheral metabolic function, emphasizing important sex differences in biomarker outcomes. Our results highlight a potential mechanism by which APOE4 alters cytokine levels in the hippocampus, which may create an environment promoting the formation of AD pathology. Our newfound correlations between peripheral metabolic markers and cytokine signatures based on *APOE* genotype suggest a valuable opportunity for future AD biomarker development.

## 2. Methods

### 2.1. Experimental Design and Statistical Analysis

The study objectives were (1) to determine how aging-related cytokine signatures in the hippocampus are affected differently by APOE3 and APOE4, (2) identify whether cytokine levels can predict systemic metabolic outcomes, and (3) uncover genotype-specific differences in the relationship between brain cytokine levels and systemic metabolism. We separated humanized APOE3 and APOE4 mice into eight study groups, each with n = 8: (1) 3-month-old APOE3 females, (2) 3-month-old APOE4 females, (3) 18-month-old APOE3 females, (4) 18-month-old APOE4 females, (5) 3-month-old APOE3 males, (6) 3-month-old APOE4 males, (7) 18-month-old APOE3 males, and (8) 18-month-old APOE4 males. This sample size was determined based on our power analysis estimated from published studies of APOE4 vs. APOE3 animals(Johnson et al., 2017), indicating 8 animals is sufficient to observe an effect size of 1.2 or greater with 80% power and alpha value of 0.05.

All mice in this study were housed in the same vivarium throughout the lifespan. Starting at 8 weeks of age, mice were weighed on a weekly basis. For each animal, 24-hr food intake, insulin sensitivity, and glucose tolerance were measured at 8, 7, and 3 days before euthanasia, respectively. Mice were euthanized at the indicated age and the hippocampus was dissected and flash frozen for further analysis.

All data are expressed as mean ± standard error of the mean. Statistical analyses for metabolic data were conducted in Graph Pad Prism 9 (v 9.2.0) using 2-way ANOVA with genotype × age (adiposity, food intake, fasting glucose, area under the curve (AUC), fasting insulin), 2-way repeated measures ANOVA genotype × time (glucose tolerance test (GTT) and insulin tolerance test (ITT) curves), mixed-effects analysis genotype × time (body mass), or 3-way repeated measures ANOVA genotype × age × time (insulin concentration during GTT). Statistical significance was determined using an error probability level of p < 0.05. When appropriate, we followed ANOVA analysis with Tukey’s or Šidák’s multiple comparisons test. We performed ROUT and Grubbs’ outlier test to identify and remove outliers (none found). Regressions comparing adiposity to glucose levels were conducted in R and visualized using ggplot2.

### 2.2. Animals

All procedures were approved by the Penn State College of Medicine Institutional Animal Care and Use Committee (PROTO201800531). A heterozygous APOE3 breeding pair (B6.Cg-Apoe^em2(APOE*)Adiuj^/J) was purchased from Jackson Laboratory, Bar Harbor, ME, USA, and bred to homozygosity, with homozygosity confirmed by qPCR. A homozygous APOE4 breeding pair (B6(SJL)-Apoe^tm1.1(APOE*4)Adiuj^/J) was also purchased from Jackson Laboratory. Mice were fed a standard chow diet (Teklad 2018, Envigo) with *ad libitum* access to food and water. Animals were maintained on a 12 hr light/12 hr dark cycle, with lights on at 07:00 and lights off at 19:00. Starting at 8 weeks of age, mice were weighed on a weekly basis. Weighing began at 14:00 every Monday and mice were weighed in the same order and on the same scale each week. Weighing at the same time and interval is important as mouse weights fluctuate with circadian rhythm(Kawamura, 2020; Minematsu et al., 1991). Food intake per cage was measured by weighing the food over a 24-hr time period. Each mouse’s individual food intake was calculated by dividing the difference in food weight over 24 hr by the number of mice, followed by dividing each individual mouse’s food intake by their body mass for a g/g food intake.

### 2.3. Insulin and Glucose Tolerance Testing

Whole body insulin and glucose action were assessed in conscious mice. For each animal, we conducted an ITT and GTT at 7 days and 3 days before euthanasia, respectively. For the ITT, mice were fasted for 4 hr and then injected intraperitoneally with insulin (0.75 U/kg of regular U-100 insulin diluted in saline, Novolin). A tail vein blood sample was taken at baseline and at 15, 30, 60, 90, and 120 min post-injection to measure blood glucose levels with a glucometer (Accu-Chek Performa, Roche). For the GTT, mice were fasted for 4 hr and then injected intraperitoneally with 50% dextrose (2 g/kg, Hospira). A tail vein blood sample was taken at baseline and at 15, 30, 60, 90, and 120 min post-injection to measure blood glucose levels. An additional blood sample was taken at baseline, 15, and 120 min with a micro-hematocrit capillary tube (Fisher) for measurement of plasma insulin concentration. Due to circadian rhythm fluctuations in glucose levels(Tingley et al., 2021), all mice began fasting at 08:30, with protocol beginning at 12:30 for all GTT and ITT assessments. Given potential differences in baseline fasting glucose among groups, changes in blood glucose during ITT and GTT procedures were normalized to baseline levels and summarized as area under the curve (AUC). Plasma insulin was measured using a mouse ultrasensitive enzyme-linked immunosorbent assay (ELISA) (ALPCO) according to the manufacturer’s protocol. Samples were run in duplicate and values below the detection limit were assigned the detection limit value of 0.115 ng/mL.

### 2.4. Body Composition

The morning of euthanasia, body mass and composition (grams of fluid, fat, and lean mass) were measured in conscious mice using a Bruker Minispec LF50 quantitative nuclear magnetic resonance analyzer (Billerica), according to the manufacturer’s protocol.

### 2.5. Euthanasia and Tissue Collection

Mice were euthanized via cervical dislocation without anesthesia. The brain was removed immediately following death and placed in dissection media (Hanks’ Balanced Salt solution buffered with 11 mM HEPES) and hemisected with a razor blade. The left hemisphere was put in 10% phosphate buffered formalin (Fisher) at 4°C and right hemisphere was dissected to remove the hippocampus. The hippocampus was homogenized in a 300 uL solution of protease inhibitor cocktail (Millipore Sigma) diluted 100x in RIPA buffer (Boston BioProducts #P8340). Hippocampal homogenate was incubated at 4°C for at least 20 min, then centrifuged at 5,000 g for 5 min. Supernatants were transferred to fresh tubes, snap-frozen in liquid nitrogen, and stored at −80°C until cytokine analysis.

### 2.6. APOE Concentration

APOE concentration in hippocampal tissue homogenate was measured using an APOE ELISA kit (Sigma Aldrich), according to manufacturer’s instructions. The assay range is 1.64 ng/mL – 400 ng/mL, thus samples were diluted accordingly with previous literature (Gee et al., 2006; Riddell et al., 2008; Sullivan et al., 2011; Ulrich et al., 2013; Wahrle et al., 2004).

### 2.7. Immunohistochemistry

The left hemisphere was passively fixed in 10% phosphate buffered formalin at 4°C for a week, then moved to a 0.1 M phosphate buffered saline (PBS) solution, pH 7.4, at 4°C for 1-4 weeks, before embedding in paraffin. Brains were then sliced sagittally with thickness of 5 μm onto charged slides and dried overnight. Slides were washed thrice with xylene, twice with 100% ethanol, then once each with 90% ethanol, 70% ethanol, and ddH_2_O for 5 min each. Slides were then boiled for 15 min with 0.01 M citrate buffer (pH 6.0), washed with PBS, 3% hydrogen peroxide, and PBS on an orbital shaker at slow speed for 5 min each. Slides were blocked with 10% normal goat serum for 20 min, then washed with PBS. Primary antibodies for glial fibrillary acid protein (GFAP; Invitrogen PA110019) and ionized calcium binding adaptor molecule 1 (IBA1; Invitrogen MA527726) were added at 1:1000 and incubated at 4°C overnight in a humidity chamber. The next day, slides were washed thrice with PBS, followed by 1 hr incubation with secondary antibodies (Goat anti-mouse Alexa Fluor 555 (Invitrogen A21425) and Goat anti-rabbit Alexa Fluor 488 (Invitrogen A11070)) in a humidity chamber. We then washed slides thrice with PBS, then twice each with 95% ethanol, 100% ethanol, and xylene prior to mounting. Slides were imaged using an Olympus BX63 fluorescence microscope, with fixed scaling across all images. Images were analyzed for GFAP and IBA1 density using ImageJ (NIH). Hippocampal cells were counted by a blinded scientist using the cell counts plugin, and cell density was calculated by dividing by total hippocampal area.

### 2.8. Cytokine Concentration

Total protein concentration of each sample was quantified using a Pierce™ BCA Protein Assay (Thermo Scientific). For Luminex analysis, samples were thawed on ice and diluted in PBS to a final protein content of 40 ug per well, which we have found to be sufficient for concentrations of most cytokines to fall in the linear range of the assay when measured in the brain tissue of this mouse model. Protein concentrations of 32 cytokines were assessed on the Luminex FLEXMAP 3D platform using a MILLIPLEX Mouse Cytokine/Chemokine Magnetic Bead Panel (MCYTOMAG-70K) according to the manufacturer’s protocol, with accommodation for a 384-well plate format: magnetic beads and antibodies were each applied at half volume. All samples were assayed in technical triplicate.

We constructed standard curves and interpolated sample cytokine concentrations using a five-parameter logistic regression(Cardillo, 2021) in MATLAB (version R2019b). We prepared the interpolated sample cytokine concentration data for analysis using an automated in-house cleaning pipeline, available for download from GitHub at https://github.com/elizabethproctor/Luminex-Data-Cleaning. We first removed all observations with bead count below 15 and calculated pairwise differences between each of the three technical replicates. If any replicate was separated from the other two by greater than twice the distance between those two, we designated it an outlier and removed it from further analysis (2% of measurements). When all 3 wells of a triplicate of sample wells recorded <15 beads, we took the average of the wells with greater than 7 beads (only 1% of measurements). We then calculated the average of the remaining technical replicates for use in downstream analysis. Cytokines with >75% of sample concentrations outside the bounds of extrapolation were removed (to minimize noise).

### 2.9. Partial Least Squares Discriminant Analysis

Data were mean-centered and unit-variance scaled in preparation for performing partial least squares (PLS) analysis. Cytokine concentrations were included as the X-block (predictors) and genotype, age, glucose tolerance, or adiposity as the Y-block (output). PLS was performed in R (v.3.6.1) using ropls (v.1.16.0)(Thévenot et al., 2015) and visualized using ggplot2 (v.3.2.1)(Villanueva and Chen, 2019). We performed cross-validation with one-third of the data and calculated model confidence by comparing our model’s predictive accuracy of cross-validation to the distribution of cross-validation accuracies of 100 randomized models. All references to “model accuracy” refer to the cross-validation accuracy. Randomized models were constructed by randomly permuting class assignment to the preserved X-block, which conserves the data landscape and provides a true control.

All PLS models were orthogonalized on the first latent variable (LV1), so that variance in parameters most correlated with the groupings of interest was maximally projected onto LV1. For each model, we chose the number of latent variables that minimized cross-validation error, and maintained this number of latent variables when constructing the distribution of randomized models. We calculated variable importance in projection (VIP) scores to quantify the contribution of each cytokine to the prediction accuracy of the overall model. VIP scores were calculated by averaging the weight of each cytokine on every latent variable across the entire model, normalized by the percent variance explained by each respective latent variable:

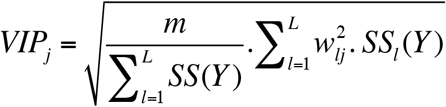

where *m* is the total number of predictors, *l* is the latent variable, *L* is the number of latent variables in the model, *w*_*lj*_ is the weight (inverse loading) of predictor *j* on latent variable *l*, and *SS*_*l*_*(Y)* is the variation in class Y explained by latent variable *l*. We note that, as a consequence of normalization, the average of squared VIP scores is 1, thus VIP > 1 is our criterion for a parameter having greater than average contribution to the model.

## 3. Results

### 3.1. APOE genotype, age, and sex modify hippocampal cytokine levels

APOE4 has been associated with inflammation in the brain of AD patients(Friedberg et al., 2020) and mouse models(Litvinchuk et al., 2021; Rodriguez et al., 2014; Shi et al., 2017) in the presence of amyloid and tau pathology. To determine whether APOE4 promotes an inflammatory environment in the brain over the course of aging independent of amyloid and neurofibrillary proteinopathies, we quantified cytokine protein concentrations in the hippocampus of young (3 months) and aged (18 months) APOE4 and APOE3 knock-in mice. These ages in mice are approximately equivalent to ages 20 and 60 years in humans(Dutta and Sengupta, 2016), allowing us to study the differences between young adulthood, when no molecular-level disease-related changes have been detected, and the prodromal period before clinical AD onset but when the initiating molecular-level processes driving AD are expected to begin.

Since aging is the greatest risk factor for AD, we first asked how the aging process changes cytokine levels in the hippocampus, which plays an important part in memory and is one of the earliest and most severely affected brain regions in AD, and how these age-related signaling changes differ depending on *APOE* genotype. We employed the supervised machine learning tool Partial Least Squares (PLS), which enables us to identify interdependent changes in a set of measured variables as they relate to our outcome or grouping of choice. Immune signaling is highly complex, with many interacting pathways, and thus individual cytokine levels are not independent of one another. PLS uses linear combinations of variables (here, cytokine concentrations) to predict membership in defined groups (here, *APOE* genotype and age)(Geladi and Kowalski, 1986; Wold et al., 1984), allowing us to quantify the complex changes in overall signaling states and networks that are lost when performing univariate analysis. These “signatures” of cytokine changes help us identify more subtle shifts in signaling that have predictive, and therefore potential mechanistic, value for defining the APOE4-induced environment in the brain that promotes AD risk.

As expected, we were able to easily distinguish young from aged mice in each genotype and sex based on cytokine levels (**Fig. 1A-1D**), indicating that cytokine signaling in the brain changes with aging. The cytokines that contributed most highly to the predictive accuracy of our model (as assessed by VIP score, Methods) in APOE3 female mice were upregulation of LIX, IP-10 (CXCL10), IL-2, and IL-1α with downregulation of IL-10 and GM-CSF (**Fig. 1A**). In APOE4 female mice, VIPs were increased levels of MIG and IP-10 but decreased levels of MIP2, MIP-1α, IL-1β, IL-13, and IFNγ (**Fig. 1C**). Male mice had their own unique signatures, where aged APOE3 male mice increased hippocampal production of IP-10, IL-2, IL-1α, IL-17, IL-15, and IL12 while downregulating eotaxin (CCL11) (**Fig. 1B**), while APOE4 males increased production of IP-10, IL-2, IL-12, and INFγ while downregulating MIP-1α in older age (**Fig. 1D**). The increase in IP-10 seen across all genotypes and sexes speaks to a partially common denominator in the known phenomenon of “inflammaging,” the increase in low-grade, generalized inflammation with aging(Bharath et al., 2020; Franceschi et al., 2018).

**Figure 1.**
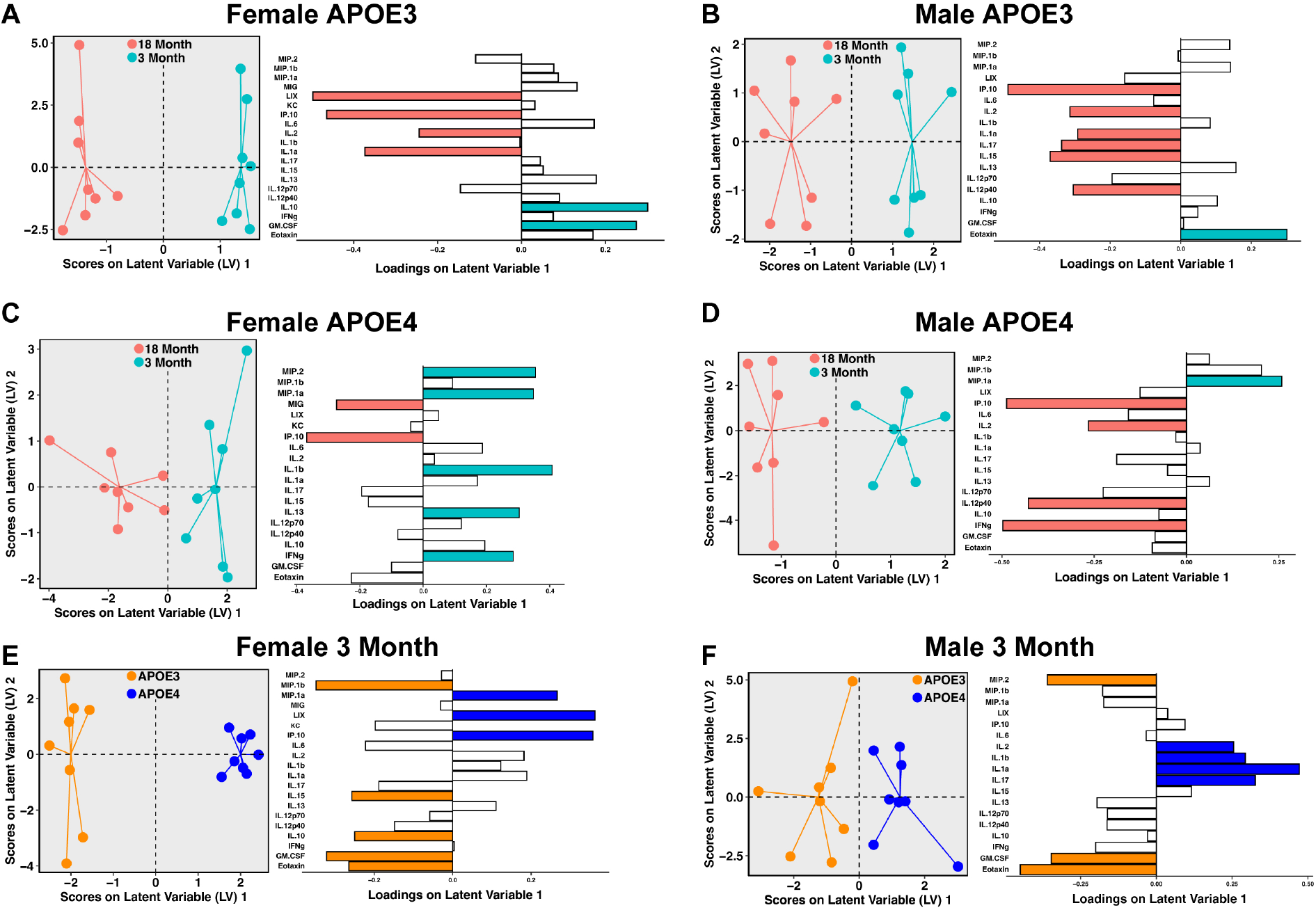
Cytokine levels increase in the hippocampus of aging APOE3 and APOE4 mice. Scores plot and loadings for Partial Least Squares Discriminant Analysis (PLSDA) of APOE3 **A**) female (5 LV, accuracy 91.67%, confidence 99.99%) and **B**) male hippocampal cytokines separated by age (4 LV, accuracy 75.89%, confidence 98.19%). Scores plot and loadings for PLSDA of APOE4 **C**) female (2 LV, accuracy 71.01%, confidence 96.87%) and **D**) male hippocampal cytokines separated by age (4 LV, accuracy 66.56%, confidence 89.82%). Scores plot and loadings for PLSDA of 3-month **E**) female (5 LV, accuracy 86.00%, confidence 99.97%) and **F**) male hippocampal cytokines separated by genotype (2 LV, accuracy 66.67%, confidence 86.29%). For PLSDA models A-D, pink and teal loadings are negative and positive variable importance in projection scores (VIPs) >1, respectively, where pink cytokines are elevated in 18-month mice and teal cytokines are elevated in 3 month mice. For PLSDA models E and F, orange and blue loadings are negative and positive VIPs >1, respectively, where orange cytokines are elevated in APOE3 mice and teal cytokines are elevated in APOE4 mice. *n* = 8 for all groups.

### 3.2. APOE4 alters cytokine protein levels differently in young male and female mice

Next, we asked how cytokine levels are affected by *APOE* genotype at young and old ages of male and female mice. Using PLS, we found that in young female mice, *APOE4* genotype was most strongly defined by higher levels of MIP-1α, LIX (CXCL5), and IP-10, with lower levels of MIP-1β, IL-15, IL-10, GM-CSF, and eotaxin in the hippocampus (**Fig. 1E**). In contrast, we found that in young male mice, *APOE4* genotype was most strongly defined by upregulation of IL-2, IL-1β, IL-1α, and IL-17, with downregulation of MIP-2, GM-CSF, and eotaxin in the hippocampus (**Fig. 1F**).

In 18-month old animals of both sexes, PLS models for predicting genotype based on hippocampal cytokines were non-significant (female prediction accuracy 54%, confidence 59%; male prediction accuracy 47%, confidence 6.8%), indicating that the cytokine signatures of APOE3 and APOE4 mice are not distinctly different at old age. Taken together, we found that hippocampal cytokine secretion signatures change during the aging process in both genotypes, and that at young age, APOE3 and APOE4 mice of each sex are easily distinguishable by their cytokine profiles but converge in old age.

### 3.3. APOE3 and APOE4 mice do not differ in gliosis at young nor old age

Since decreases in MIP-1α were observed in male and female APOE4 animals over the lifespan, we next determined whether these genotypic differences were supported by decreases in activation of astrocytes and microglia. We measured gliosis in the hippocampus using immunohistochemistry and found no significant differences between genotypes in microglial or astrocytic cell count or activation at either age (**Fig. 2A-H**). However, the number of GFAP-positive astrocytes decreased between 3 and 18 months of age in APOE3 mice (**Fig. 2A**), telling an opposite story from the easily-discernible genotype-specific patterns of cytokine protein signatures we saw in aged mice, evidencing that cytokine signatures are an additional level of detail needed to study immune function and activity in the brain.

**Figure 2.**
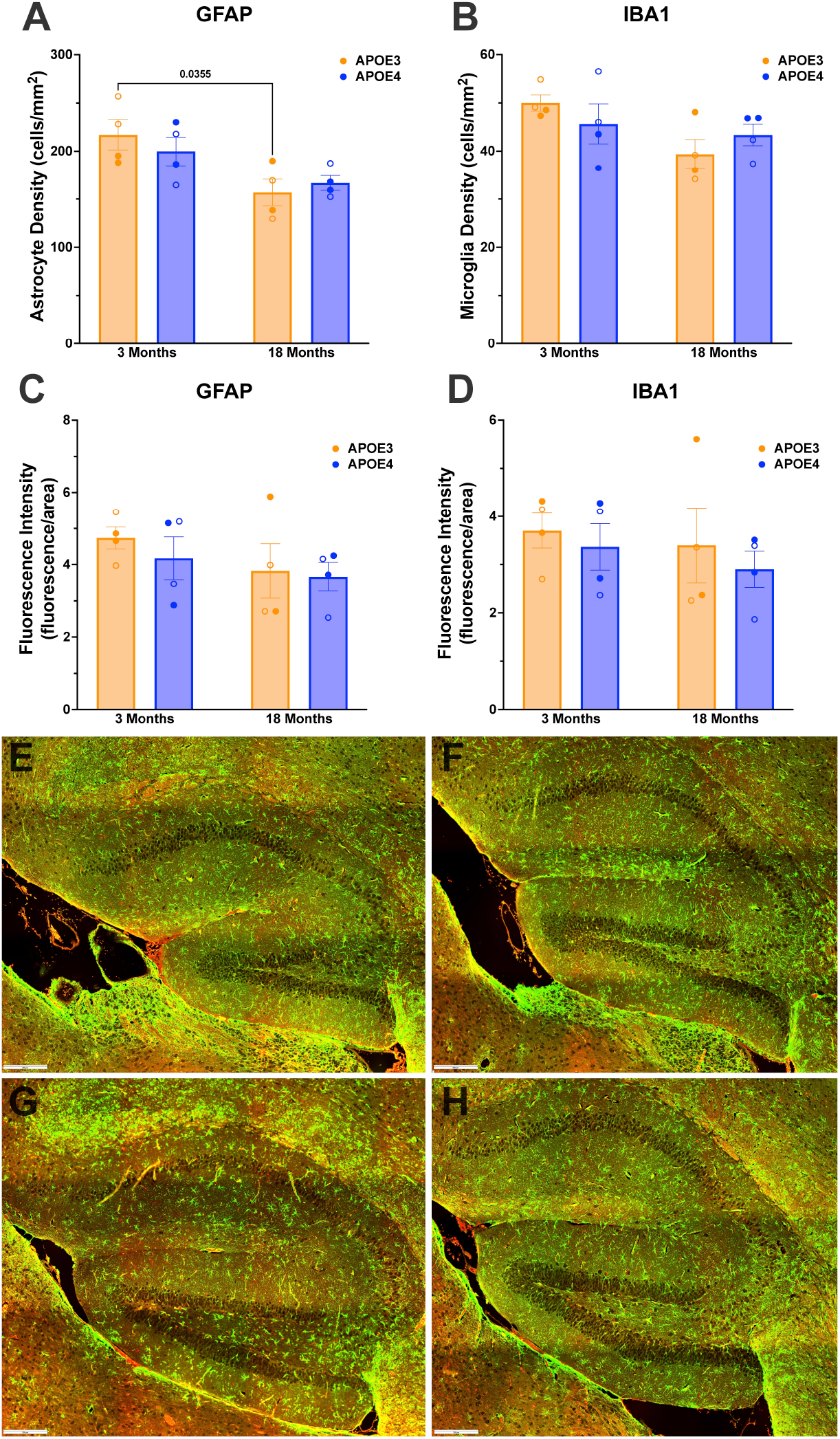
*APOE* genotype does not differentially affect gliosis in young or old mice. Representative quantification (**A**-**D**) of immunohistochemistry stains showcasing gliosis determined by GFAP (astrocytes) and IBA1 (microglia) in **E**) APOE3 3-month, **F**) APOE4 3-month, **G**) APOE3 18-month, and **H**) APOE4 18-month within hippocampal brain slices. Scale bar, 200 μm. Main effects of cell counts **A**): *P*_*age*_ *=* 0.005, *P*_*genotype*_ *=* 0.788, interaction *P =* 0.331) and **B**) *P*_*age*_ *=* 0.049, *P*_*genotype*_ *=* 0.956, interaction *P =* 0.182). Main effects of fluorescence intensity **C**) *P*_*age*_ *=* 0.211, *P*_*genotype*_ *=* 0.516, interaction *P =* 0.714) and **D**) *P*_*age*_ *=* 0.477, *P*_*genotype*_ *=* 0.443, interaction *P =* 0.884). Data are mean +/-SEM and were analyzed by 2-way-ANOVA with Tukey’s test when significant, *n* = 4 mice per group (mixed sex). All *P* values on graphs reflect post-hoc analyses. Open circles are males, closed circles are females.

### 3.4. Female APOE4 mice experience less age-related weight gain and adiposity than their APOE3 counterparts

Previous studies link systemic metabolism with hippocampal changes(Seto et al., 1983; Tingley et al., 2021); and while best known for its function as a lipid transporter, *APOE* genotype impacts both lipid and glucose metabolism. Thus, to measure systemic metabolic changes due to APOE4, and link these changes to the brain, we first weighed mice weekly and determined body composition at 3 and 18 months of age. APOE3 and APOE4 mice gained weight at a steady rate across the lifespan (**Fig. 3A and 3B**). While we did not observe significant differences in weight between genotypes at any individual time point, mixed-effects models proved a statistically significant attenuation of weight gain over time in female mice, with APOE4 female mice decreasing their rate of weight gain compared to APOE3 female mice starting at approximately age 50 weeks (**Fig. 3A**).

**Figure 3.**
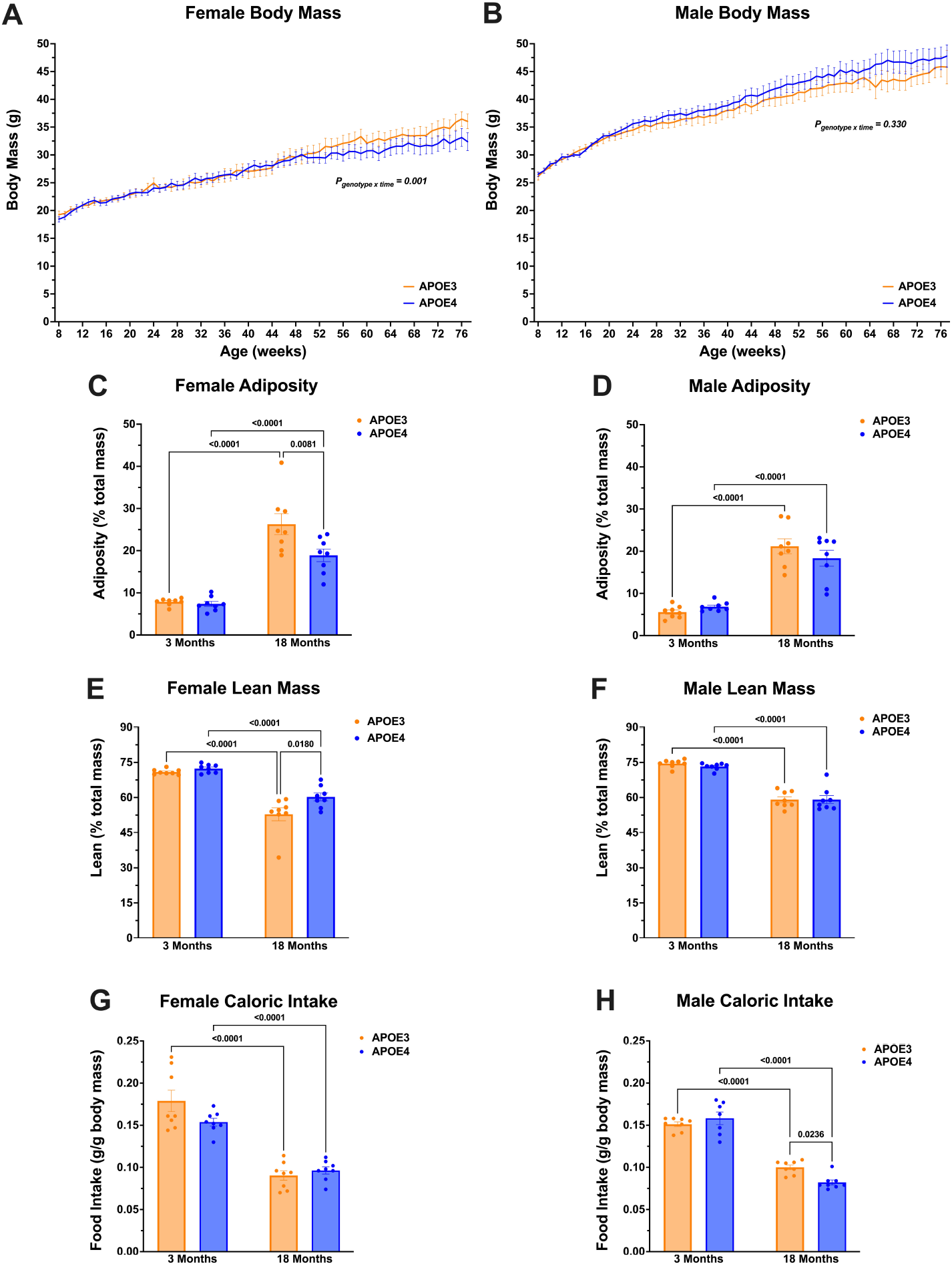
APOE4 mice experience less aging-related increases in adiposity than their APOE3 counterparts in female but not male mice. Weekly body mass of 18-month-old mice from 8 weeks to 77 weeks of age in **A**) females (fixed effects: *P*_*time*_< 0.001; *P*_*genotype*_ = 0.496; interaction *P* = 0.001) and **B**) males (fixed effects: *P*_*time*_< 0.001; *P*_*genotype*_ = 0.311; interaction *P* = 0.330). No individual time points were significantly different between APOE3 and APOE4 in female or male mice. Body fat percentage (adiposity) of **C**) female (main effects: *P*_*age*_ *<* 0.001, *P*_*genotype*_ *=* 0.014, interaction *P =* 0.028) or **D**) male (main effects: *P*_*age*_ *<* 0.001, *P*_*genotype*_ *=* 0.562, interaction *P =* 0.133) mice measured by qNMR. Lean body percentage of **E**) female (main effects: *P*_*age*_ *<* 0.001, *P*_*genotype*_ *=* 0.012, interaction *P =* 0.081) or **F**) male (main effects: *P*_*age*_ *<* 0.001, *P*_*genotype*_ *=* 0.587, interaction *P =* 0.572) mice measured by qNMR. Food intake of **G**) female (main effects: *P*_*age*_ *<* 0.001, *P*_*genotype*_ = 0.217, interaction *P* = 0.049) or **H**) male (main effects: *P*_*age*_ *<* 0.001, *P*_*genotype*_ *=* 0.207, interaction *P =* 0.006) mice, normalized to per gram of body weight. Data are mean +/-SEM and were analyzed by (A and B) Mixed-effects model (REML) followed by Šidák’s multiple comparison test; (C-H) 2-way-ANOVA followed by Tukey’s test. *n* = 8 for all groups. All *P* values on graphs reflect post-hoc analyses.

To understand the physiological distribution of age-related weight gain, we used qNMR to quantify lean, fat, and fluid mass of each mouse. We found that while APOE3 and APOE4 mice had similar body composition at a young age, aged APOE4 female mice have significantly lower body fat mass (about 40% lower) than their APOE3 counterparts (**Fig. 3C**), paired with a 14% increase in lean mass (**Fig. 3E**). The attenuated weight gain and lower body fat in APOE4 female mice were not a result of decreased food intake; we quantified the crude weight of food consumed by mice in each cage as grams consumed per gram of body weight (**Fig. 3G**) and, while aged mice of both genotypes consumed less food per gram of body weight than their young counterparts (as expected from previous studies of food consumption in aging(Morley, 2001; Pilgrim et al., 2015)), food consumption was not significantly different between APOE3 and APOE4 female mice at either age. In male mice, no differences existed between APOE3 and APOE4 counterparts in terms of body mass, adiposity, and lean mass (**Fig. 3D** and **3F**), although APOE4 males ate slightly less (18%) food per gram of body weight than their APOE3 counterparts (**Fig. 3H**).

### 3.5. APOE4 protects against aging-related decreases in glucose tolerance in female mice

The differences we observed between APOE3 and APOE4 female mice in age-related weight and adipose gain, despite similar food consumption, suggests a difference in how these animals metabolize food. Given that the chow that the animals were fed is very low in fat and consists predominantly of carbohydrate (17% fat, 60% carbohydrate, 23% protein), a systemic alteration in glucose metabolism is a potential mechanism (Martínez-Martínez et al., 2020). Fasting glucose is a measure of how well the body maintains glucose homeostasis in the absence of food, demonstrating the ability to regulate glucose, insulin, and glucagon levels. We measured fasting blood glucose levels in 3-and 18-month-old animals of both genotypes and found no significant differences between female genotypes at either age, nor did fasting blood glucose change with aging for either genotype (**Fig. 4A**). We also did not observe any genotypic differences in fasting glucose levels of male mice at young or old age; however, APOE4 male mice had increased fasting glucose levels at 18-months of age, compared to 3-month counterparts (**Fig. 4B**).

**Figure 4.**
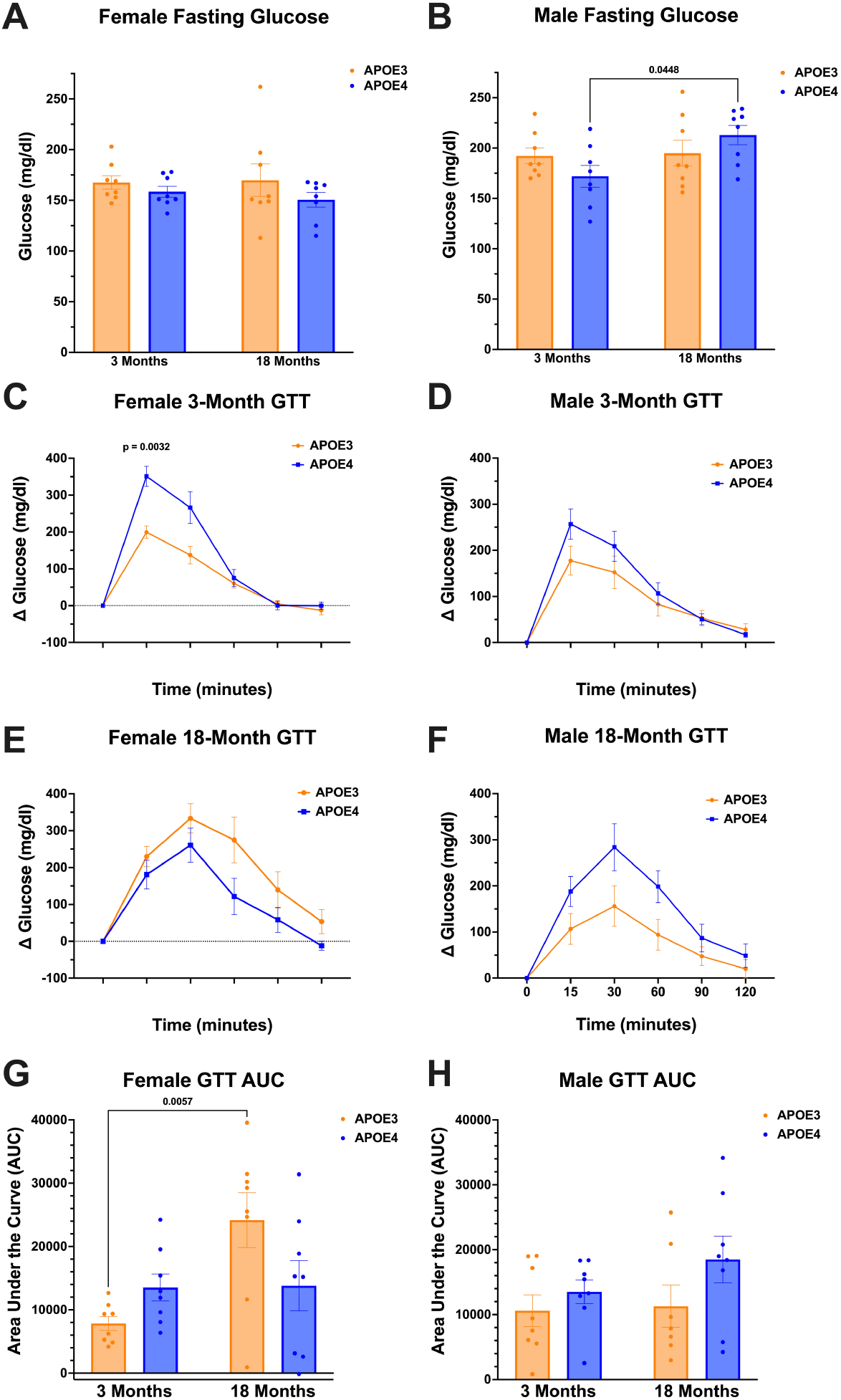
APOE4 protects against aging-related decreases in glucose tolerance independent of glucose-stimulated insulin secretion in female but not male mice. Fasting glucose levels of 3- and 18-month APOE3 and APOE4 **A**) female (main effects: *P*_*age*_ *=* 0.212, *P*_*genotype*_ = 0.309, interaction *P* = 0.845) and **B**) male (main effects: *P*_*age*_ *=* 0.046, *P*_*genotype*_ = 0.910, interaction *P* = 0.076) mice. Glucose tolerance test (GTT) of 3-month-old APOE3 and APOE4 **C**) female (*P*_*genotype*_ *=* 0.012) and **D**) male (*P*_*genotype*_ *=* 0.296) mice. GTT of 18-month-old APOE3 and APOE4 **E**) female (*P*_*genotype*_ *=* 0.096) and **F**) male (*P*_*genotype*_ *=* 0.093) mice. GTT AUC comparison across age and genotype for **G**) female (*P*_*genotype*_ *=* 0.465, *P*_*age*_ *=* 0.014, interaction *P =* 0.017) and **H**) male (*P*_*genotype*_ *=* 0.086, *P*_*age*_ *=* 0.328, interaction *P =* 0.460) mice. Data are mean +/-SEM and were analyzed by (A, B, G, and H) 2-way-ANOVA followed by Tukey’s test when appropriate; (C-F) 2-way repeated measures ANOVA, followed by Šidák’s multiple comparison test when significant. *n* = 7-8 for all groups. All *P* values on graphs reflect post-hoc analyses.

While fasting glucose did not vary between APOE4 and APOE3 animals, differences in weight gain and adiposity despite similar food consumption may be caused by decreased ability to metabolize glucose. A glucose tolerance test (GTT) uses a glucose injection to measure the body’s endogenous insulin function, where high GTT area under the curve (AUC) values suggest glucose intolerance. At 3 months of age, APOE4 female mice exhibited significantly higher 15-min peak blood glucose concentrations compared to APOE3 female mice but did not show a significantly different GTT AUC overall, indicating similar levels of glucose tolerance (**Fig. 4C and 4G**). In contrast, aged APOE3 female mice experienced a significant decrease in their ability to handle large glucose loads, while APOE4 female mice were protected from these aging-related decreases in glucose tolerance as measured by the overall GTT AUC (**Fig. 4E** and **4G**). On the other hand, male APOE4 mice did not exhibit different GTT responses than APOE3 male mice at young nor old age (**Fig. 4D, 4F**, and **4H**).

### 3.6. Insulin sensitivity decreases in male but not female mice at 18 months of age

Since APOE4 female mice did not experience the aging-related decline in glucose tolerance seen in APOE3 mice, we next asked whether endogenous insulin action could explain this difference. To determine the insulin sensitivity of APOE3 and APOE4 mice, we quantified glucose uptake in response to a measured insulin bolus using an insulin tolerance test (ITT), where a more negative ITT AUC value indicates better insulin sensitivity. Insulin sensitivity was not significantly different between APOE3 and APOE4 female mice in either age group (**Fig. 5A and 5B**), nor between young and aged mice of either genotype (**Fig. 5C**).

**Figure 5.**
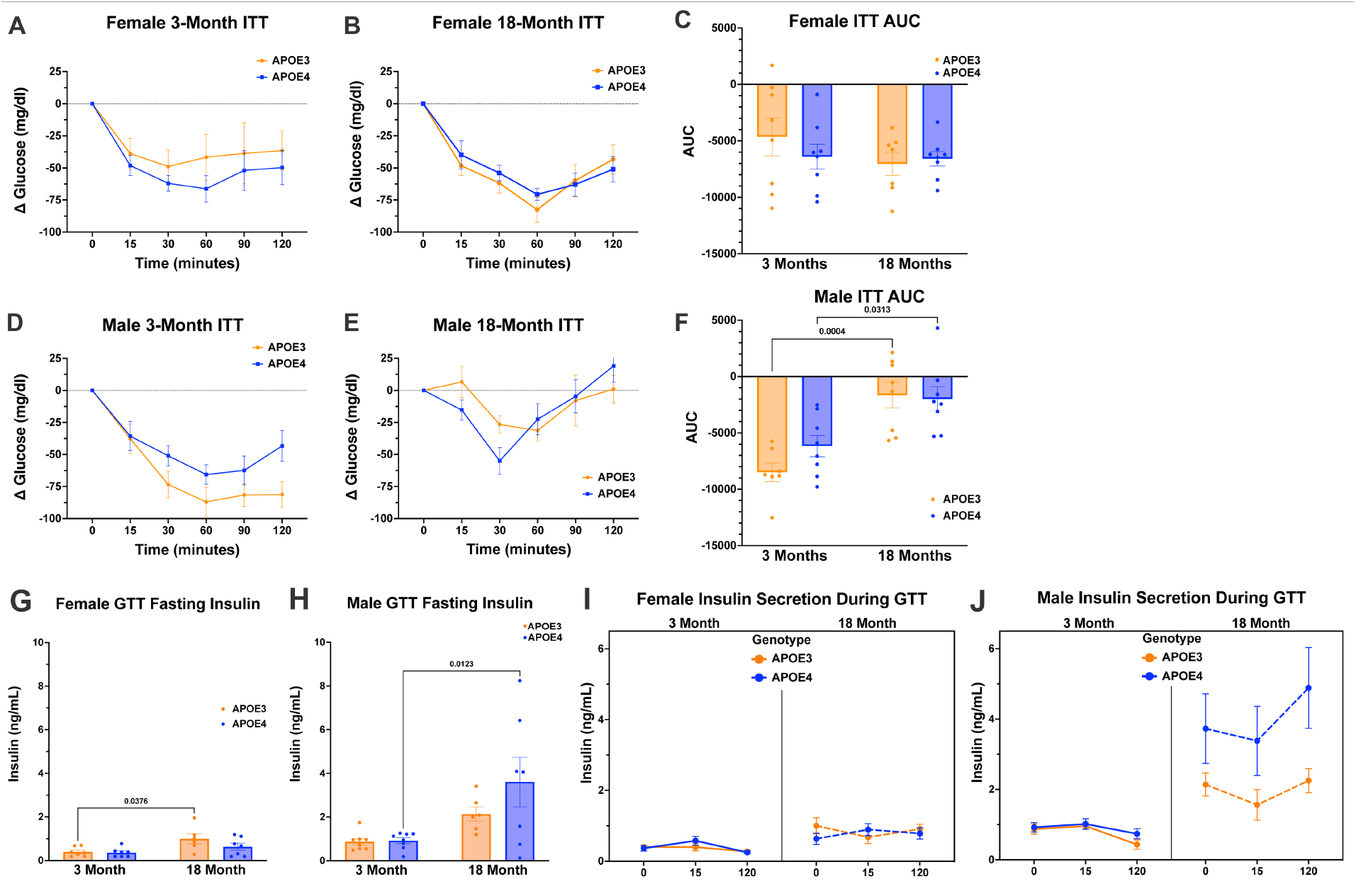
Aging increases insulin levels of APOE4 male but not female mice. Insulin tolerance test (ITT) of 3-month-old APOE3 and APOE4 **A**) female (*P*_*genotype*_ *=* 0.389) and **D**) male (*P*_*genotype*_ *=* 0.091) mice. ITT of 18-month-old APOE3 and APOE4 **B**) female (*P*_*genotype*_ *=* 0.739) and **E**) male (*P*_*genotype*_ *=* 0.770) mice. Area under the curve (AUC) analysis of APOE3 and APOE4 ITT at both 3 and 18 months in **C**) female (*P*_*age*_ *=* 0.286, *P*_*genotype*_ = 0.582, interaction *P =* 0.346) and **F**) male (*P*_*age*_ *=* < 0.001, *P*_*genotype*_ = 0.339, interaction *P =* 0.203) mice. Fasting insulin concentration in **G**) female (*P*_*age*_ *=* 0.002, *P*_*genotype*_ *=* 0.099, interaction *P =* 0.367) and **H**) male (*P*_*age*_ *=* 0.002, *P*_*genotype*_ *=* 0.198, interaction *P =* 0.229) mice. Insulin concentration at 0-, 15-, and 120-min post glucose injection of **I**) female (*P*_*age*_ < 0.001, *P*_*genotype*_ = 0.659, *P*_*time*_ = 0.477, *P*_*time x age*_ *=* 0.124, *P*_*time x genotype*_ *=* 0.012, *P*_*genotype x age*_ *=* 0.676, *P*_*genotype x age x time*_ *=* 0.593) and **J**) male (*P*_*age*_ = 0.002, *P*_*genotype*_ = 0.110, *P*_*time*_ = 0.527, *P*_*time x age*_ *=* 0.002, *P*_*time x genotype*_ *=* 0.747, *P*_*genotype x age*_ *=* 0.180, *P*_*genotype x age x time*_ *=* 0.818) mice. Data are mean +/-SEM and were analyzed by (A, B, D, and E) 2-way repeated measures ANOVA, (C, F, G, and H) 2-way-ANOVA with Tukey’s test when significant, with *n* = 7-8 for all groups. I and J were analyzed by 3-way REML, with n = 5-8 per time/group. All *P* values in figure plots reflect post-hoc analyses.

We next sought to identify whether a lack of endogenous insulin production was the cause of decreased glucose tolerance in aged APOE3 female mice. We measured plasma insulin concentration at fasted baseline and at 15 and 120 min after glucose injection. We found no difference in fasting insulin levels between APOE3 and APOE4 females at either age, but an increase in APOE3 18-month-old females, compared to their 3-month counterparts (**Fig. 5G**). These higher fasting insulin levels in APOE3 aged females, mirror a similar phenomenon in humans where aging increases fasting insulin levels(Johnson et al., 2010).

To clarify whether endogenous insulin is properly released in response to spikes in blood glucose, we measured insulin levels at the 15 and 120 min time points after glucose injection. If the insulin response is working properly, insulin levels should spike to allow glucose into the cells, then diminish to baseline concentration at 120 min. Across ages and genotypes in female mice, the insulin secretion across the GTT did not change (**Fig. 5I**). This insignificant change in insulin measured over time likely indicates that the insulin level spike in response to glucose and returned to baseline between 0-15 min and thus was not observed at our timepoints.

Overall, these results suggest that although female APOE4 and APOE3 mice maintain normal baseline glucose levels, insulin sensitivity, and insulin secretion during aging, APOE4 female mice diverge during aging from APOE3 in their glucose tolerance. Thus, aging impairs glucose tolerance in APOE3, but not APOE4 female mice via a mechanism independent of insulin sensitivity or glucose-stimulated insulin secretion.

In male mice, ITT AUC was not different between *APOE* genotypes at either age but decreased substantially in both APOE3 and APOE4 mice between 3 and 18 months of age, showing that aged male mice have decreased insulin sensitivity(**Fig. 5D, 5E**, and **5F**). Similar to females, we saw no genotypic differences in fasting insulin at either age in males. Instead, we found an increase in APOE4 fasting insulin levels with age and no change in APOE3 fasting insulin levels with age (**Fig. 5H**). Whereas on average the fasting insulin levels doubled during aging in females, fasting insulin tripled in males. The increases in fasting insulin levels during aging in males also increased in variability, especially in the APOE4 animals. Also similar to females, across ages and genotypes in male mice, the insulin secretion across the GTT did not change (**Fig. 5J**), indicating that the insulin response to glucose likely spiked between measured timepoints.

These results suggest that dissimilarly to female mice, male APOE3 and APOE4 mice are similar at young and old ages, yet aging has large, significant effects on insulin secretion and sensitivity, indicating that while aging affects insulin production and usage in males, these metabolic parameters are not affected by *APOE* genotype.

### 3.7. Hippocampal cytokine levels predict adiposity of APOE3 and APOE4 mice

A remaining gap in the AD field is how APOE-related systemic effects impact the brain (and vice versa). To investigate a potential mechanism for the relationship between systemic metabolism and hippocampal cytokine protein expression patterns, we performed PLS regression (PLSR), which is similar to our previous PLS analysis except that instead of separating animals based on discrete categories (i.e., genotype or age group), latent variables predict a measurement with a continuous numeric value (e.g., adiposity, GTT AUC, ITT AUC). Our goal was to determine whether specific cytokine signatures are predictively associated with certain systemic metabolic outcomes, and in doing so uncover genotype-specific differences in the relationship between brain cytokine expression and systemic metabolism.

First, we focused on each *APOE* genotype separately to ask which changes in cytokine signaling within the hippocampus correlate with body adiposity. We constructed hippocampal cytokine signatures capable of predicting body fat percentage in mice of all ages within 5-7% of the measured value upon cross-validation (**Fig. 6A-6D**). This predictive accuracy was highly statistically significant, in all cases resulting in >99% model confidence compared to random chance, indicating that we can reliably predict systemic metabolic outcomes based on hippocampal cytokine levels in male and female mice of each genotype.

**Figure 6.**
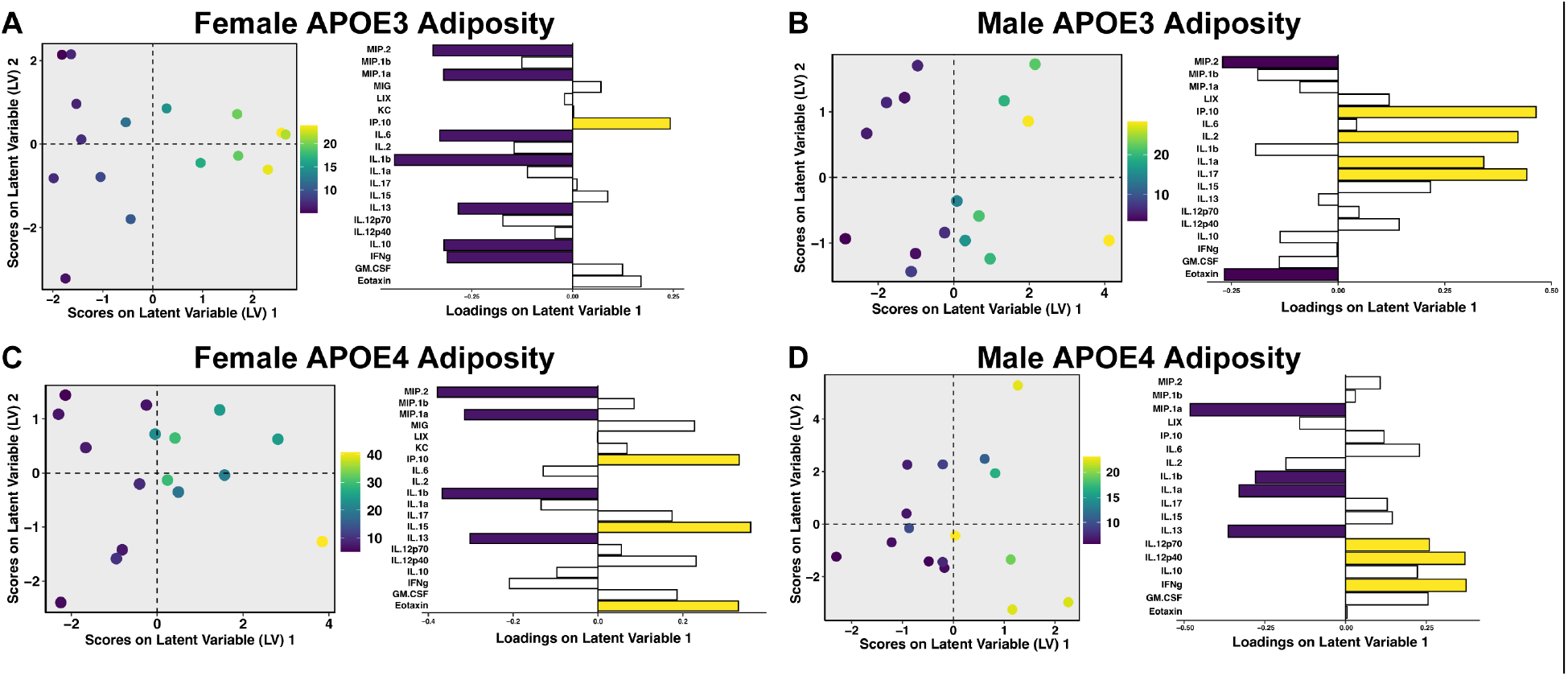
Hippocampal cytokine signaling predicts adiposity of aging female and male mice. Scores plot and loadings for PLS regression (PLSR) of APOE3 **A**) female (4 LV, RMSECV 6.66, confidence 99.98%) and **B**) male hippocampal cytokines separated by adiposity (1 LV, RMSECV 5.31, confidence 100%). Scores plot and loadings for PLSR of APOE4 **C**) female (5 LV, RMSECV 4.63, confidence 99.96%) and **D**) male hippocampal cytokines separated by adiposity (1 LV, RMSECV 6.79, confidence 99.99%). Violet and yellow loadings are negative and positive variable importance in projection (VIPs) >1, respectively. *n* = 8 for all groups.

We saw that as APOE3 female mice increase in adiposity, they have lower levels of MIP-2, MIP-1α, IL-6, IL-1β, IL-13, IL-10, and INFγ, with increased production of IP-10 (CXCL10) (**Fig. 6A**). Conversely, in APOE4 females, MIP-2, MIP-1α, IL-1β, IL-13 all decrease, while IP-10, IL-15, and eotaxin increase, with increasing adiposity (**Fig. 6C**). APOE3 and APOE4 female mice share similar signatures with MIP-2, MIP-1α, IL-1β, and IL-13 decreasing, while IP-10 increases with adiposity. Major differences between APOE3 and APOE4 females highlight that APOE4 females secrete more IL-15 and eotaxin when mice have higher adiposity while APOE3 females decrease several cytokines as they gain adiposity. These signatures indicate that the relationship between hippocampal cytokine levels and body adiposity is fundamentally altered in APOE4 versus APOE3 female mice, with APOE4 mice showing stronger immune activation with increased adiposity, evidenced by increased levels of IL-15 and eotaxin, two inflammatory cytokines.

As APOE3 male mice increase in adiposity, they have lower levels of MIP-2 and eotaxin, and increased production of IP-10, IL-2, IL-1α, and IL-17 (**Fig. 6B**). Conversely, in APOE4 males, MIP-1α, IL-1α, IL-1β, and IL-13 decrease, while IL-12 and INFγ increase as adiposity increases (**Fig. 6D**). In contrast to females, males of *APOE3* and *APOE4* genotype share no similarities in their cytokine profiles when correlating adiposity with hippocampal cytokine protein expression. These male signatures support the conclusion that the relationship between peripheral adiposity and hippocampal cytokine levels are altered by *APOE* genotype, with APOE4 male mice activating dissimilar cytokines to APOE3 male mice.

### 3.8. Hippocampal cytokine levels predict glucose tolerance of APOE3 and APOE4 mice

Since we observed significant genotype differences in glucose tolerance of females and differing genotype-dependent predictive relationships between adiposity and hippocampal cytokine signaling, we next performed PLS analysis to regress hippocampal cytokine levels against GTT AUC measurements from young and old male and female APOE3 and APOE4 mice, to determine whether cytokine levels in the brain can predict systemic glucose tolerance.

Similar to adiposity, each of our four models had high predictive accuracy and model confidence (here all >99%). We found that as glucose tolerance decreases (GTT AUC increases) in APOE4 female mice, hippocampal levels of several measured cytokines decreased, with decreases in MIG, LIX, IP-10 (CXCL10), IL-17, and IL-12 being most predictive of glucose tolerance (**Fig. 7C**). In APOE3 female mice, IL-10 production decreased with lower glucose tolerance, while MIP-2, LIX, IP-10, IL-2, IL-1α, and IL-12 increased (**Fig. 7A**). The opposing effects in APOE4 versus APOE3 mice, especially manifested in LIX, IL-12, and IP-10 signaling, suggests that with decreasing glucose tolerance, APOE3 female mice experience upregulation of canonically pro-inflammatory cytokines while APOE4 females experience a muted cytokine secretion response.

**Figure 7.**
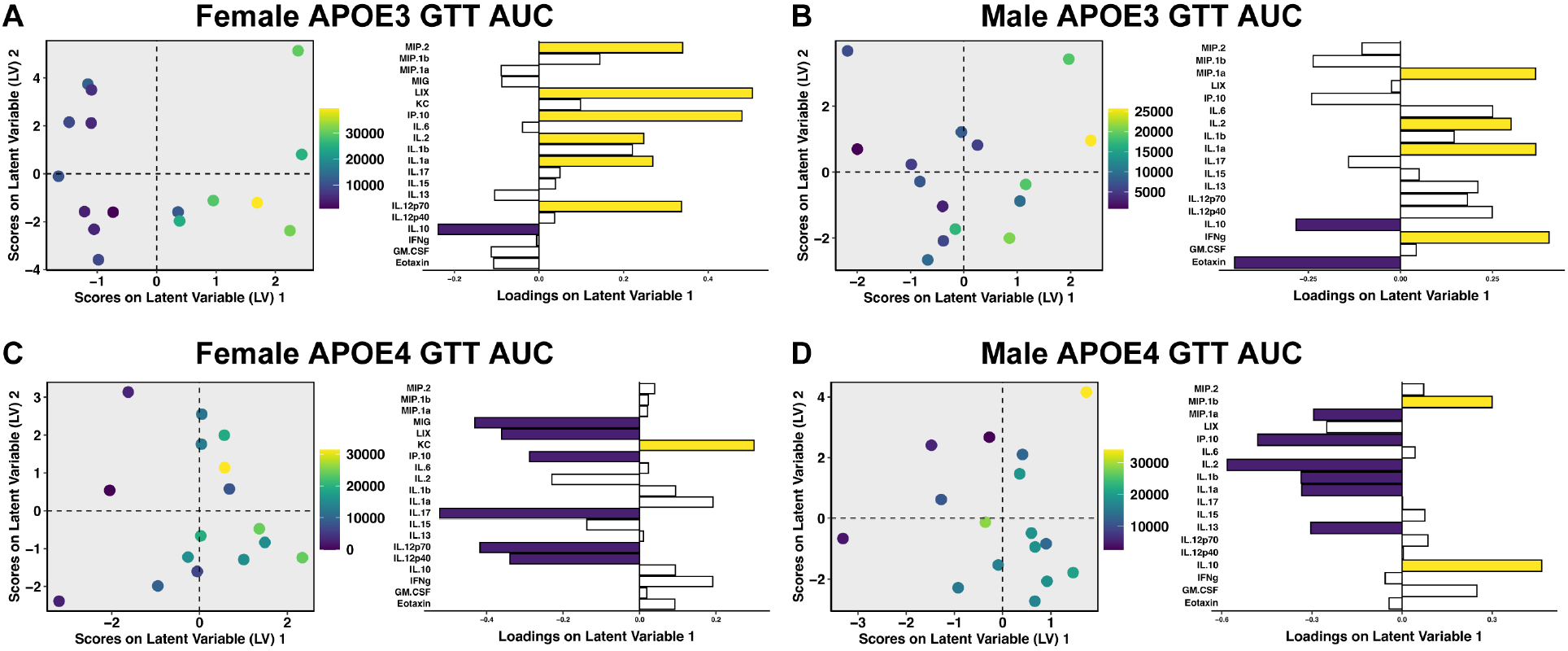
Hippocampal cytokine signaling predicts glucose tolerance of aging female and male mice. Scores plot and loadings for PLSR of APOE3 **A**) female (2 LV, RMSECV 9,672.463, confidence 99.99%) and **B**) male hippocampal cytokines separated by glucose tolerance (GTT AUC) (1 LV, RMSECV 6,755.54, confidence 100%). Scores plot and loadings for PLSR of APOE4 **C**) female (1 LV, RMSECV 9,273.22, confidence 99.85%) and **D**) male hippocampal cytokines separated by glucose tolerance (1 LV, RMSECV 6,592.83, confidence 100%). Violet and yellow loadings are negative and positive variable importance in projection (VIPs) >1, respectively. *n* = 8 for all groups.

APOE4 males upregulated release of hippocampal MIP-1β and IL-10 as glucose tolerance decreased, while downregulating MIP-1α, IP-10, IL-2, IL-1β, IL-1α, and IL-13 (**Fig. 7D**). In contrast, APOE3 male mice upregulated MIP-1α, IL-2, IL-1α, and IFNγ as glucose intolerance decreased, and downregulated IL-10 and eotaxin (**Fig. 7B**). Like female mice, males show opposing *APOE* genotypic differences, especially manifested in IL-10, MIP-1α, IL-2, and IL-1α signaling. Additionally, male signatures are unique from the females and further highlight sex and *APOE* genotype as inflammation-defining parameters.

### 3.9. Hippocampal cytokine levels predict insulin sensitivity of APOE3 and APOE4 mice

Since we observed significant genotype differences in insulin sensitivity and production with aging in male mice, we next asked whether hippocampal cytokine levels could predict insulin sensitivity. Using PLSR we regressed hippocampal cytokine concentrations against ITT AUC values from young and old male and female APOE3 and APOE4 mice, and again found major differences between genotypes and sexes. APOE3 female mice decrease LIX and IL-12, and increase MIP-1α, MIG, IL-6, IL-17, and IL-13, as their insulin sensitivity decreases (**Fig. 8A**). Conversely, APOE4 female mice decrease IL-10 and increase MIP-1α, IL-1α, IL-13, and IFNγ as their insulin sensitivity decreases (**Fig. 8C**). In male mice, APOE3 carriers decreased eotaxin and increased IP-10, IL-2, IL-1α, Il-17, IL-15, and IL-12 as their insulin sensitivity decreased (**Fig. 8B**). APOE4 males increased levels of the largest range of hippocampal cytokines, MIP-2, IL-6, IL-17, IL-12, IFNγ, GM-CSF, and eotaxin, downregulating none, as they lost insulin sensitivity (**Fig. 8D**). Overall, these results suggest that cytokine secretion increases in the hippocampus as insulin sensitivity decreases and that the specific patterns of cytokine expression are defined by sex and *APOE* genotype.

**Figure 8.**
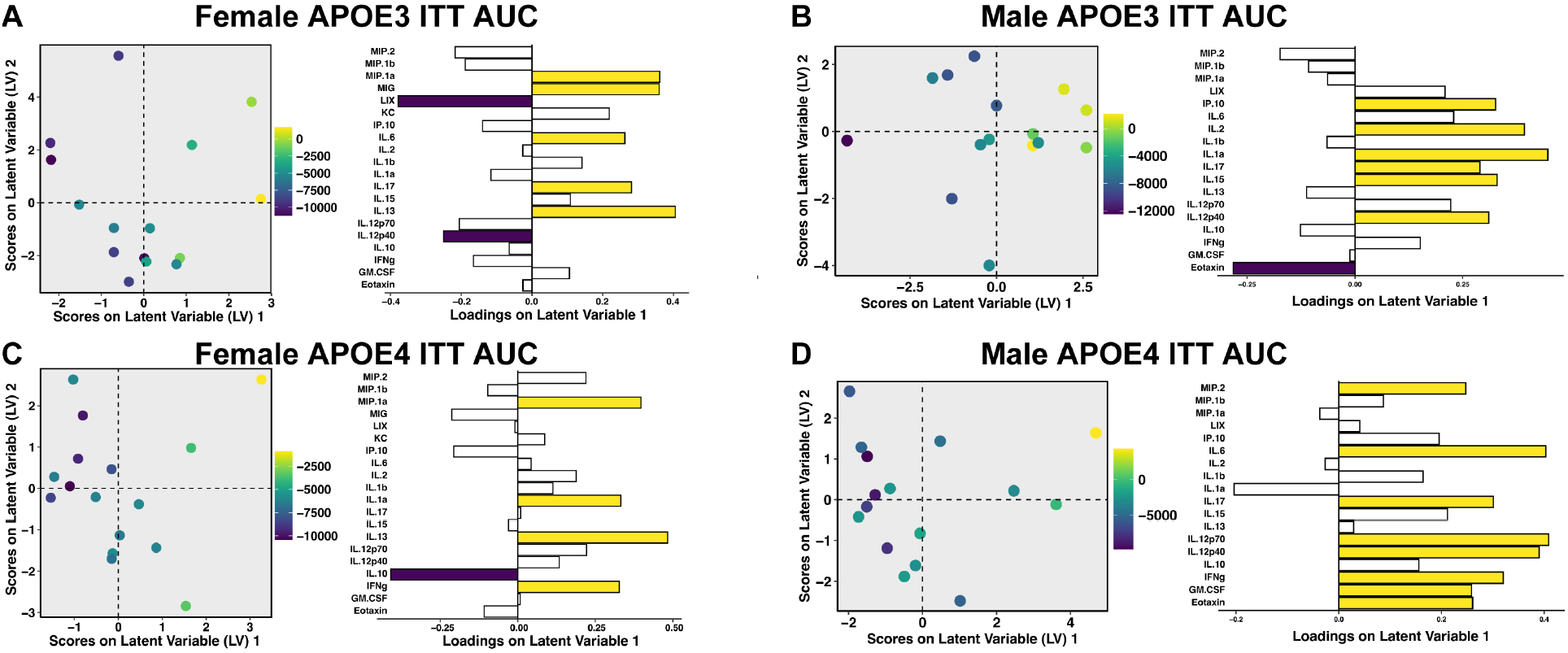
Hippocampal cytokine signaling predicts insulin sensitivity of aging female and male mice. Scores plot and loadings for PLSR of APOE3 **A**) female (2 LV, RMSECV 3,774.48, confidence 99.98%) and **B**) male hippocampal cytokines separated by insulin tolerance (ITT AUC) (1 LV, RMSECV 2,772.86, confidence 100%). Scores plot and loadings for PLSR of APOE4 **C**) female (1 LV, RMSECV 2,530.54, confidence 99.97%) and **D**) male hippocampal cytokines separated by ITT AUC (1 LV, RMSECV 3,491.79, confidence 99.97%). Violet and yellow loadings are negative and positive variable importance in projection (VIPs) >1, respectively. *n* = 8 for all groups.

### 3.10. APOE genotype, not hippocampal concentration, impacts metabolic parameters in male and female mice

To better understand which factors impact the metabolic parameters we measured, we performed several regression analyses. In both males and females, glucose and adiposity were not correlated in APOE3 or APOE4 carriers (**Fig. 9A** and **9B**), suggesting that fasting glucose level and adiposity are two independent variables of peripheral metabolism for APOE3 and APOE4 mice.

**Figure 9.**
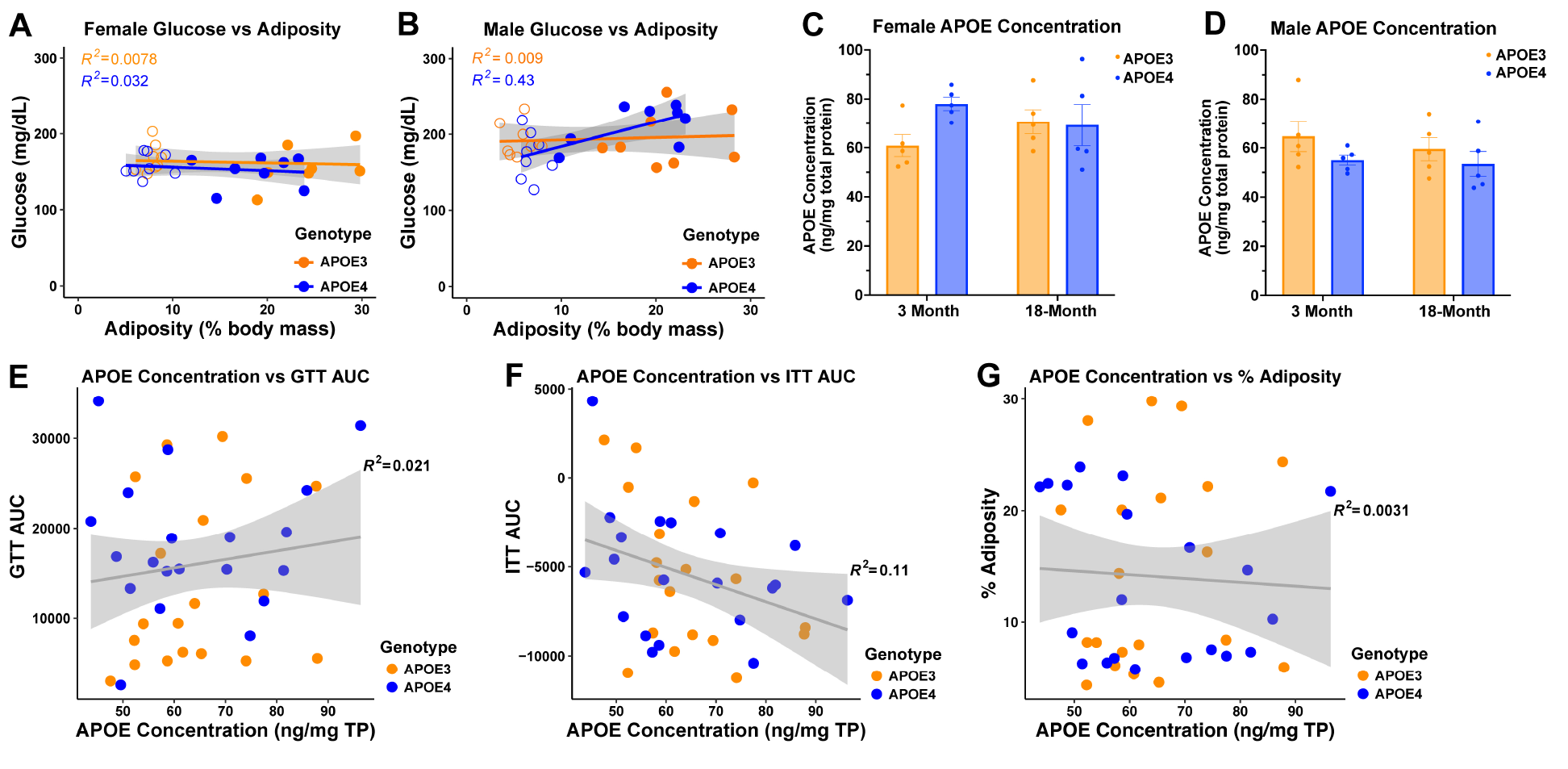
APOE concentration in the hippocampus does not correlate with metabolic parameters. **A**) Female and **B**) male linear regressions of adiposity versus fasting glucose in APOE3 and APOE4 mice, where open circles are 3-month-old and filled circles are 18-month-old mice. Concentration of APOE in **C**) female and **D**) male APOE3 and APOE4 mice. Regressions of mixed male/female, young/old hippocampal APOE concentration with **E**) GTT AUC, **F**) ITT AUC and **G**) adiposity. Data in (C and D) are mean +/-SEM and were analyzed by 2-way-ANOVA with Tukey’s test when significant, with *n* = 5-8 for all groups. Data in (A, B, E, F, G) were plotted in R with regression line and R^2^ values calculated with *lm* function; grey area shows 95% confidence region, with *n* = 7-8 for all groups.

To further identify why differences in metabolic outcomes were different in APOE3 and APOE4 mice, we next measured the concentration of APOE in the hippocampus of male and female APOE3 and APOE4 mice. We found that there were no differences due to age of genotype in male or female mice (**Fig. 9C** and **8D**). We next calculated whether APOE concentration correlated with the peripheral metabolic parameters we measured and found that APOE concentration was not correlated with glucose tolerance, insulin sensitivity, or adiposity (**Fig. 9E-9G**), suggesting that *APOE* genotype and not concentration, was the culprit in our cytokine level changes.

## 4. Discussion

APOE4 is both the strongest and the most common genetic risk factor for AD, but the impact of APOE4 in promoting a disease environment outside of its interaction with AD protein pathology is incompletely understood. APOE4 is known to disrupt systemic metabolism(Martínez-Martínez et al., 2020; Torres-Perez et al., 2016), while dysfunctional systemic metabolism is known to increase risk of AD(Martínez-Martínez et al., 2020; Razay et al., 2007), and APOE4 is known to drive AD proteinopathies(Castellano et al., 2011; Huynh et al., 2017; Litvinchuk et al., 2021; Liu et al., 2017; Shi et al., 2017), yet these three variables are persistently examined and analyzed separately. Studying brain and systemic pathology separately is disadvantageous because the effects of APOE on systemic metabolism and brain immune signaling are interdependent. Ignoring this relationship discounts potentially important avenues for therapeutic or preventative interventions for APOE4-implicated conditions. To identify APOE4-related systemic metabolic changes that co-vary with altered cytokine protein profiles in the brain, we quantified the relationships between functional measures of systemic metabolism and a broad sample of hippocampal cytokine signaling pathways in young and old humanized APOE4 and APOE3 knock-in mice. Importantly, this work was performed in the absence of tau and amyloid-β pathologies in the brain, highlighting the importance of the disease-promoting environment created by APOE4 prior to, and independent of, the development of proteinopathies. We have demonstrated that APOE3 and APOE4 mice differ in their peripheral metabolism and brain cytokine levels, and that the relationships between peripheral metabolism and brain immune systems differ distinctly by *APOE* genotype and sex.

While best known for its function as a lipid transporter, APOE impacts both lipid and glucose metabolism. In the brain, APOE4 decreases insulin signaling and glucose metabolism(Ong et al., 2014; Perkins et al., 2016). In the periphery, evidence in humans is mixed, where APOE4 has been shown to decrease glucose tolerance, increase insulin resistance, or have no effect(Martínez-Martínez et al., 2020). Although APOE itself cannot cross the blood-brain barrier, systemic metabolic changes due to *APOE* genotype control levels of a host of other immune- and metabolism-altering molecules that do have this ability and can thus have profound effects in the brain. We found that cytokine levels in the hippocampus can predict adiposity, glucose tolerance, and insulin sensitivity with high accuracy. A new study connected peripheral liver-expressed APOE4 to decreased cognition and synaptic plasticity in the brains of mice(Golden and Johnson, 2022; Liu et al., 2022). Previous studies have also identified a link between systemic metabolism and the hippocampus, where electrophysiological pulses (sharp wave-ripples) in the hippocampus regulate peripheral glucose concentration(Tingley et al., 2021). Communication between the hippocampus and systemic metabolism may involve autonomic signaling from the hippocampus to the pancreas and liver, or through hippocampal signaling to the hypothalamus(Tingley et al., 2021), a major metabolic control center in the brain. Additionally, electrical stimulation of the hippocampus has been shown to regulate systemic glucose and insulin levels(Lathe, 2001; Seto et al., 1983). These aspects of hippocampal function led us to question whether the connection we found between hippocampal cytokine signatures and systemic metabolism is uni-directional or bi-directional. In its proposed role as a controller of glucose metabolism, altered hippocampal immune signaling could be the stimulus for *APOE* variant-specific differences in adiposity and glucose tolerance. Alternatively, peripheral changes in adiposity, glucose tolerance, and insulin sensitivity may impact hippocampal cytokine levels through upregulation of factors that can pass the blood-brain barrier, or through affecting vascular integrity and allowing infiltration of peripheral immune cells and signaling molecules. Finally, and perhaps most likely given the evidence for both aspects of directionality, both of these scenarios could be occurring simultaneously. Future experiments are needed to determine mechanisms for the sex differences we observed here, as well as the method and directionality of communication between these co-varying pathways.

Human *APOE4* carriers are known to have an increased risk for metabolic syndrome, a risk coupled to body fatness(Torres-Perez et al., 2016). However, when fed a low-fat diet, we found that APOE4 female mice experienced less weight and fat gain with age than did APOE3 mice. Therefore, in the absence of pre-existing obesity, APOE4 female mice are, in contrast from APOE3 mice, protected from aging-related impairments in glucose tolerance and insulin sensitivity. Indeed, previous studies have shown that APOE4 has a protective effect on weight gain compared to APOE3 in humans and mice, even in scenarios of high fat diet(Arbones-Mainar et al., 2016, 2008; Segev et al., 2016; Tejedor et al., 2014). Despite these protective effects, incidence of AD is significantly higher in *APOE4* carriers, suggesting that the AD risk conferred by APOE4 is a separate mechanism from risk due to metabolic syndrome.

Importantly, our findings highlight the importance of considering cytokine profiles as a highly connected network, allowing for interaction and covariation in the levels of the various cytokines assayed. Without this multivariate approach, we would not have been able to identify meaningful signatures of alteration to the brain’s immune state, nor to define the quantitative relationship of these signatures with systemic metabolism. The specific APOE4 cytokine signatures we found, and their corresponding systemic metabolic changes suggest that APOE4 could have additional AD risk effects beyond its ability to increase amyloid-β(Liu et al., 2017) and tau(Shi et al., 2017) aggregation and its ability to reduce clearance of these aggregates(Castellano et al., 2011). Both male and female APOE4 animals exhibited significant, model-defining decreases in MIP-1α over the lifespan. MIP-1α is a chemokine secreted by cells to attract microglia, the brain’s macrophage cell type, which scavenge and clear plaques and damaged synapses to protect and maintain neuron health. Significant loss of MIP-1α, which we did not observe in APOE3 animals, could suggest decreased ability to clear disease insults from the brain due to lack of microglia recruitment, potentially resulting in a loss of neuronal resilience. In females, the contrasting cytokine signatures we observed in the hippocampus of APOE4 versus APOE3 mice may alter hippocampal signaling to the hypothalamus, ameliorating the aging-related weight gain, increased adiposity, decreased glucose tolerance, and increased baseline insulin secretion observed in their APOE3 counterparts. While previous transcriptomic studies in the brain have found that age has a greater effect on gene expression than does *APOE* genotype(Zhao et al., 2020), we conversely found that protein levels are highly dependent on *APOE* genotype, highlighting the contribution of the various levels of regulation between genotype and phenotype. We found that protein levels of IL-17, IL-6, and IFNγ increased in APOE3 and APOE4 males and females as insulin sensitivity decreased. Similarly, T-helper 17 (Th17) cells, which can infiltrate the brain and produce high levels of IL-17, IL-6, and IFNγ, are upregulated in insulin resistance and type 1 and type 2 diabetes(Ip et al., 2016; Nicholas et al., 2019; Tesmer et al., 2008; Zhang et al., 2019). Future studies evaluating cytokine protein levels in the hypothalamus of APOE knock-in animals will allow a better understanding of direct effects on the brain’s metabolic control center.

Across their lifespan, APOE3 female mice gained more weight and body fat than APOE4 female mice. This increased body fat was independent of caloric intake in relation to body weight. One limitation of our study is that we did not quantify the type and location of fat mass. Importantly, subcutaneous, visceral, and brown fat each have different metabolic value and function: visceral fat is associated with cardiovascular and metabolic impairments, while brown fat is known to be metabolically active and beneficial(Antonopoulos and Tousoulis, 2017). Indeed, previous studies have shown that APOE4 mice have increased fatty-acid oxidation and brown fat mass(Arbones-Mainar et al., 2016), thus the increased fat that we measured in APOE3 mice is likely visceral fat(Arbones-Mainar et al., 2016). This possibility is further supported by the decreased glucose tolerance and raised fasting insulin levels that we observed in aged APOE3 mice, but additional studies are needed to confirm this hypothesis.

We found that APOE4 females are protected from the age-associated decreases in glucose tolerance experienced by their APOE3 counterparts. The lack of change in insulin sensitivity and insulin secretion response during GTT in all female groups, in combination with the increased fasting insulin levels in aged female APOE3 mice, suggest that the mechanism of APOE4 protection against aging-related decrease in glucose tolerance in females is insulin-independent. However, observation of glucose levels at finer time resolution and earlier than the 15 min GTT time point will clarify whether any earlier changes occur in the peak and shape of the insulin spike. Of note, higher doses of glucose administered to mice with higher body mass may influence GTT results due to calculation based on total mass, but our results likely remain similar to dosing based on lean mass as previously seen in comparison of dosing methods in obese mice(Jørgensen et al., 2017). Our results suggest that males of both genotypes, as well as APOE3 female mice, tend toward hyperinsulinemia with aging, similar to effects seen in humans(Janssen, 2021). Young APOE3 females were the most glucose tolerant, while old APOE3 mice had the lowest tolerance. APOE4 animals of both sexes maintained glucose tolerance across the lifespan but experienced a shift in their peak glucose level timing. Other potential mechanisms contributing to the diminished aging-related decrease in glucose tolerance in APOE4 mice include altered intestinal glucose absorption(Ussar et al., 2017), glucose effectiveness (the ability of glucose to regulate its production by the liver and global utilization)(Hu et al., 2021), expression and activity of glucose transporters (GLUTs)(Chadt and Al-Hasani, 2020), and changes in circulating metabolites(Chadt and Al-Hasani, 2020; Williams et al., 2016).

While the differences in glucose tolerance and fasting insulin levels between APOE4 and APOE3 female mice were not statistically significant at 18 months of age, aged APOE4 mice appeared to be moving toward lower GTT AUC and lower fasting insulin levels, suggesting that extending the current experiment out to 24 months may result in observation of larger, statistically significant differences between *APOE* genotypes. However, the 18-month-old mice used in this study are approximately equivalent to humans at age 60. Thus, extending the study to include older animals may change the implications and relevance for AD risk to a study of AD onset and progression. Notably, we studied mice in the context of healthy aging, in the absence of AD proteinopathies, and thus our results are important for other aging-related diseases including cardiovascular disease, dyslipidemia, stroke, and other dementias. In the present study, we profiled both male and female mice but did not directly compare them as their results are so qualitatively different. The current AD research environment is moving towards personalized medicine, as the cancer field has, where sex is a profound variable that must be considered for drawing conclusions on disease state and treatment plan. Specifically, females display heightened levels of significant immune dysregulation experienced by women in aging and disease, a phenomenon observed both in female AD patients(Guo et al., 2022) and in the absence of AD(Angum et al., 2020; Klein and Flanagan, 2016). Additionally, nearly two-thirds of dementia-related deaths are female(Oh and Rabins, 2019), and APOE4-related risk for AD is stronger in females(Bretsky et al., 1999; Farrer et al., 1997). Future studies should compare females and males directly to discern whether female metabolic and hippocampal cytokine signaling differ in respect to male counterparts.

## Supporting information

Hippocampal cytokine levels (pg per mL)

## Acknowledgements

This work was supported by R01AG072513 from the National Institute on Aging (EAP), R01AA209403 from the National Institute on Alcohol Abuse and Alcoholism (NAC), and start-up funds from the Penn State College of Medicine Departments of Neurosurgery and Pharmacology (EAP). RMF is supported by NIH NRSA predoctoral fellowship F31AG071131 from the National Institute on Aging. MKK and DCC are supported by training fellowship T32NS115667 from the National Institute of Neurological Disorders and Stroke. We thank Lynne Beidler for help in mouse handling and colony management and Marianne Klinger for help in brain embedding, slicing, and sample mounting. The Metabolic Phenotyping Core (RRID:SCR_022565) services and instruments used in this project were funded, in part, by the Pennsylvania State University College of Medicine via the Office of the Vice Dean of Research and Graduate Students and the Pennsylvania Department of Health using Tobacco Settlement Funds (CURE). The content is solely the responsibility of the authors and does not necessarily represent the official views of the University or College of Medicine. The Pennsylvania Department of Health specifically disclaims responsibility for any analyses, interpretations or conclusions. The Core also acknowledges support from NIH through S10OD026980.

