## Supplementary material for "Predictive link between systemic metabolism and cytokine signatures in the brain of apolipoprotein E ε4 mice": Hippocampal cytokine levels (pg per mL)

| Sample | Genotype | Sex | Age | Eotaxin | GM-CSF | IFNg | IL-1a | IL-1b | IL-2 | IL-6 | IL-10 | IL-12p40 | IL-12p70 | IL-13 | IL-15 | IL-17 | IP-10 | KC | LIX | MIG | MIP-1a | MIP-1b | MIP-2 |
|---|---|---|---|---|---|---|---|---|---|---|---|---|---|---|---|---|---|---|---|---|---|---|---|
| E3-245-H | APOE3 | Female | 3 | 38.49979986 | 22.78203826 | 37.56219055 | 108.6107888 | 20.91684787 | 23.38873254 | 11.52303574 | 59.53431477 | 4.906639778 | 31.10866613 | 518.2903684 | 17.7554531 | 18.57438524 | 41.42238419 | 12.20181509 | 138.988027 | 67.29991227 | 106.1914558 | 40.13976243 | 319.2781891 |
| E3-246-H | APOE3 | Female | 3 | 25.57151084 | 9.552209975 | 32.55289202 | 129.1525594 | 12.97946258 | 38.38422509 | 4.82933127 | 77.14852112 | 2.823899631 | 18.03555769 | 1021.129379 | 10.9430439 | 4.738393228 | 17.30980384 | 10.87091647 | 152.1511508 | 4.28102163 | 676.7624568 | 47.75114149 | 317.8260596 |
| E3-254-H | APOE3 | Female | 3 | 22.33523649 | 5.86157594 | 19.51539289 | 75.74004274 | 9.654207634 | 22.55520904 | 2.37984186 | 58.26867358 | 0.321504476 | 0 | 325.9917783 | 5.300636166 | 1.612008625 | 20.69497389 | 0 | 139.8010258 | 0 | 0 | 38.50067867 | 296.2676651 |
| E3-255-H | APOE3 | Female | 3 | 42.38762711 | 17.12958698 | 74.5189751 | 148.058206 | 12.52538768 | 44.93260069 | 3.421048115 | 76.03607479 | 19.54048529 | 17.760141 | 430.1230769 | 12.10472582 | 0 | 50.15371071 | 0 | 193.3481539 | 0 | 77.79869885 | 50.08255171 | 312.7146925 |
| E3-256-H | APOE3 | Female | 3 | 25.44777054 | 16.98443269 | 23.35518907 | 86.97988562 | 16.75345802 | 24.34538698 | 12.65088307 | 72.60421109 | 5.403408388 | 33.47315068 | 454.6960906 | 17.22782377 | 12.00031063 | 25.60343298 | 16.66462805 | 135.601974 | 0.22633621 | 0 | 50.50699254 | 298.1483974 |
| E3-257-H | APOE3 | Female | 3 | 19.87609236 | 8.120617777 | 12.44214134 | 106.0743777 | 13.20732165 | 24.56132811 | 1.978863499 | 71.55139511 | 0.089212509 | 4.807759773 | 571.3179552 | 7.438741007 | 1.329334999 | 25.90205294 | 0 | 112.3084609 | 0 | 0 | 37.00748993 | 308.217781 |
| E3-258-H | APOE3 | Female | 3 | 24.64816081 | 14.51395603 | 31.3803672 | 123.1933284 | 10.86264196 | 21.8481929 | 2.216195782 | 85.44130406 | 0.92035872 | 15.00538337 | 464.2693802 | 11.9157694 | 0.539149014 | 30.5800234 | 0 | 107.7732398 | 0 | 0 | 44.63485871 | 310.629224 |
| E3-273-H | APOE3 | Female | 3 | 18.40841434 | 5.932027259 | 12.82295734 | 39.70329282 | 3.627174636 | 5.740366613 | 0.885049072 | 62.1975865 | 0.195060897 | 9.093202858 | 267.6682558 | 2.52833451 | 0 | 18.21864205 | 0 | 131.2450335 | 0 | 0 | 38.67950181 | 307.6289219 |
| E4-225-H | APOE4 | Female | 3 | 20.71235001 | 0 | 25.66735985 | 145.1114384 | 11.03772727 | 38.24973658 | 1.573127385 | 61.70651369 | 0.784676955 | 3.767496943 | 565.1308248 | 5.15555392 | 0.957589476 | 48.42918769 | 0 | 173.9218341 | 0 | 486.6727262 | 28.54369134 | 303.2951325 |
| E4-233-H | APOE4 | Female | 3 | 21.1467899 | 0 | 31.6567027 | 126.5706581 | 19.48384661 | 44.23166195 | 1.67048812 | 66.01855364 | 1.574321194 | 16.01781571 | 769.7698282 | 5.220593026 | 1.099478152 | 40.27902756 | 0 | 262.1990724 | 3.959279226 | 1053.970343 | 33.12561023 | 297.5304431 |
| E4-235-H | APOE4 | Female | 3 | 18.55678672 | 4.308169289 | 46.40996719 | 149.7138529 | 15.81330487 | 33.16807911 | 1.216592796 | 54.41163468 | 0.690258657 | 11.1716848 | 773.5583254 | 6.237930897 | 0 | 54.59362138 | 0 | 220.6991245 | 5.006269718 | 1074.249228 | 28.81095715 | 310.3900765 |
| E4-236-H | APOE4 | Female | 3 | 21.23957712 | 5.185305 | 28.15409017 | 105.8110067 | 10.56180041 | 35.15447609 | 3.279025163 | 56.47303379 | 3.909273338 | 17.3396311 | 298.8667599 | 9.179306304 | 1.623655159 | 45.2299189 | 6.630721761 | 147.3792604 | 11.66833498 | 0 | 34.07181362 | 300.1210657 |
| E4-237-H | APOE4 | Female | 3 | 17.39717275 | 0 | 24.54855034 | 77.09416978 | 13.41759898 | 22.98283873 | 0.85087266 | 57.509338 | 0.553037499 | 6.039060874 | 319.9394938 | 2.175089352 | 0 | 41.72762591 | 0 | 217.1663049 | 0 | 299.8235295 | 29.97784408 | 307.5932522 |
| E4-238-H | APOE4 | Female | 3 | 22.06256823 | 11.3242027 | 30.57269826 | 145.8232487 | 14.09966565 | 36.652301 | 2.209838962 | 69.81718017 | 1.360012129 | 17.1244385 | 542.5616446 | 9.140472448 | 1.27269901 | 56.94113266 | 0 | 214.2279465 | 0 | 423.5105939 | 43.70233042 | 310.0727394 |
| E4-246-H | APOE4 | Female | 3 | 18.43285633 | 3.861853214 | 41.08166467 | 144.9152911 | 20.70701745 | 44.1550557 | 2.074767815 | 56.74295665 | 2.695869085 | 30.79986674 | 2079.624188 | 4.388357442 | 2.693280411 | 79.39269224 | 0.240902536 | 233.8803927 | 36.58808667 | 1601.432733 | 33.1860899 | 321.810551 |
| E4-247-H | APOE4 | Female | 3 | 18.27368338 | 4.308169289 | 19.42135888 | 108.0039589 | 11.50886987 | 19.38106582 | 1.204308266 | 67.41890945 | 0.783850562 | 9.233638468 | 235.1618435 | 3.850287499 | 0 | 52.23768294 | 0 | 193.2631925 | 0 | 119.3537494 | 35.27600702 | 313.0607258 |
| E4-8-H | APOE4 | Female | 18 | 23.74538369 | 10.28518719 | 24.15591614 | 130.6388549 | 11.12289589 | 20.51137316 | 1.898398946 | 72.17326745 | 1.88856155 | 13.05141208 | 328.6130157 | 6.963301774 | 0 | 179.5896502 | 2.118459866 | 165.2366348 | 28.00480124 | 53.00479783 | 36.55564645 | 313.9611743 |
| E4-9-H | APOE4 | Female | 18 | 21.12658547 | 3.401119568 | 37.22889439 | 120.4439126 | 15.218229 | 41.23569371 | 1.958457473 | 64.51779465 | 1.850554916 | 8.624343461 | 371.2449668 | 6.033533195 | 0 | 94.106164 | 6.055482119 | 166.8087634 | 11.06692927 | 0 | 27.13960227 | 296.2542304 |
| E4-10-H | APOE4 | Female | 18 | 26.65758498 | 6.997441302 | 28.1831016 | 116.5345765 | 4.651111432 | 27.88911198 | 1.1247079 | 54.84928505 | 1.377467984 | 0 | 308.7246217 | 8.341958888 | 0.784161848 | 91.58016684 | 5.251537406 | 136.3314112 | 0 | 0 | 38.97112269 | 297.4751698 |
| E4-22-H | APOE4 | Female | 18 | 19.81396156 | 3.037003559 | 17.59670063 | 103.0703334 | 7.796174694 | 28.83161948 | 1.060920228 | 50.86091116 | 0.168444928 | 0.353293512 | 293.5048339 | 3.91428542 | 0 | 113.6576172 | 0 | 208.3628586 | 0 | 0 | 23.45106803 | 297.5827773 |
| E4-23-H | APOE4 | Female | 18 | 20.26333522 | 3.739474282 | 10.60734949 | 120.8716599 | 9.792146341 | 23.81424704 | 0.666328475 | 53.16062042 | 1.076749613 | 7.100545265 | 136.5843 | 4.535813526 | 0 | 158.0669103 | 0 | 227.7345002 | 0 | 9.457757128 | 30.43142901 | 290.6560331 |
| E4-35-H | APOE4 | Female | 18 | 16.21359458 | 3.037003559 | 15.62151621 | 118.8254559 | 6.951559527 | 37.24803614 | 0.690898562 | 44.10619029 | 1.238840277 | 10.74489922 | 176.7646781 | 3.002367804 | 1.523917638 | 54.64785657 | 0 | 228.3179919 | 0 | 29.63981521 | 17.51014468 | 298.7690457 |
| E4-36-H | APOE4 | Female | 18 | 24.57042531 | 6.5900313 | 23.6409838 | 105.8204313 | 7.707352026 | 40.09631132 | 1.585929418 | 62.13667593 | 2.653575024 | 15.01276287 | 252.9992797 | 9.499888975 | 6.108749967 | 348.9664869 | 0 | 212.2106773 | 510.3153268 | 0 | 34.62718745 | 299.505509 |
| E4-37-H | APOE4 | Female | 18 | 22.20195811 | 4.022813024 | 20.88308214 | 100.8243145 | 6.5838487 | 31.93858271 | 0.822881671 | 56.77122932 | 3.141875822 | 26.00774177 | 7.473666129 | 10.10243678 | 0.85908461 | 88.89140353 | 0 | 237.1006035 | 0 | 0 | 36.84372305 | 292.7149228 |
| E3-30-H | APOE3 | Female | 18 | 19.37807304 | 1.358534746 | 30.78761889 | 178.4288336 | 11.42267333 | 43.68551049 | 1.411778516 | 53.08704719 | 0.484959817 | 2.196171241 | 583.3881015 | 3.143734224 | 3.415747505 | 126.4769214 | 0 | 183.0140644 | 5.736679967 | 255.7800412 | 27.13720392 | 290.0513928 |
| E3-39-H | APOE3 | Female | 18 | 30.10779965 | 21.90167851 | 43.94500205 | 176.3157418 | 26.37467863 | 50.63649476 | 8.234154561 | 64.95969727 | 9.780859376 | 40.3788071 | 687.9261526 | 22.67622596 | 15.05494756 | 101.1113418 | 18.6034905 | 283.7173666 | 2.775790623 | 171.8990171 | 61.41842055 | 330.2639643 |
| E3-50-H | APOE3 | Female | 18 | 20.61120783 | 5.565184463 | 20.17360699 | 142.3147355 | 9.163168914 | 23.52872419 | 1.977416477 | 68.15247671 | 0.820856935 | 13.75537616 | 339.2036135 | 7.434496032 | 0 | 83.4571025 | 0 | 181.8307 | 0 | 0 | 37.06086187 | 306.1074753 |
| E3-51-H | APOE3 | Female | 18 | 24.01709139 | 9.576602852 | 29.38203792 | 153.3892388 | 14.60667212 | 27.01735403 | 4.288233494 | 74.72432015 | 3.847732631 | 67.08186212 | 452.2441531 | 8.463173265 | 7.328260009 | 85.65183046 | 0 | 218.6456249 | 0 | 46.96965737 | 53.74495387 | 350.6622581 |
| E3-56-H | APOE3 | Female | 18 | 23.46854416 | 4.208516353 | 30.87237978 | 125.8552747 | 16.58226812 | 36.26157975 | 1.999593408 | 64.12549249 | 0.969129783 | 19.02344305 | 447.0683644 | 6.596760009 | 1.253704774 | 99.59330954 | 0 | 197.4750463 | 0 | 29.70075858 | 41.27299883 | 306.283116 |
| E3-57-H | APOE3 | Female | 18 | 22.248644 | 3.739474282 | 22.52317301 | 110.1965111 | 14.72888438 | 40.55410156 | 1.887661834 | 54.20823692 | 2.136242708 | 9.590494885 | 370.2947525 | 7.35867318 | 0.536317008 | 295.2076115 | 1.554171089 | 150.6709775 | 11.89694595 | 17.89570494 | 33.67457524 | 312.1933219 |
| E3-58-H | APOE3 | Female | 18 | 24.97813742 | 7.702469321 | 21.28986474 | 94.79110688 | 8.387889122 | 19.80596837 | 4.139354549 | 64.00433148 | 4.542710287 | 30.97371731 | 148.6614914 | 11.80192674 | 0.467277793 | 83.7751187 | 14.13469405 | 235.9235185 | 0 | 0 | 44.68436388 | 315.7459633 |
| E3-97-H | APOE3 | Female | 18 | 27.90157316 | 6.402445192 | 20.64296668 | 123.3545568 | 0 | 25.60231014 | 2.189774406 | 65.65194626 | 1.333020294 | 5.160098557 | 228.3141162 | 11.74307858 | 3.439716182 | 59.72489055 | 0 | 181.0378137 | 0 | 0 | 36.29882269 | 296.1104174 |

| Sample | Genotype | Sex | Age | Eotaxin | GM-CSF | IFNg | IL-1a | IL-1b | IL-2 | IL-6 | IL-10 | IL-12p40 | IL-12p70 | IL-13 | IL-15 | IL-17 | IP-10 | LIX | MIP-1a | MIP-1b | MIP-2 |
|---|---|---|---|---|---|---|---|---|---|---|---|---|---|---|---|---|---|---|---|---|---|
| E3-239-H | APOE3 | Male | 3 | 29.65449265 | 10.28518719 | 19.0231574 | 91.63914778 | 13.92434505 | 25.07072494 | 1.522640271 | 60.31583275 | 0.968032625 | 10.6850127 | 257.5096104 | 3.943794649 | 0 | 39.07372076 | 194.222273 | 24.06385182 | 33.78300515 | 326.9434852 |
| E3-240-H | APOE3 | Male | 3 | 24.28289538 | 4.35857225 | 18.97796455 | 93.46262923 | 16.73980784 | 29.6736949 | 1.826758848 | 59.81401584 | 0.785011447 | 4.801035189 | 288.7638928 | 6.344062193 | 0 | 33.35041812 | 221.8694478 | 12.82276805 | 33.03679259 | 307.5932522 |
| E3-241-H | APOE3 | Male | 3 | 30.89264217 | 5.481748725 | 23.95251806 | 143.0836464 | 15.77827958 | 22.07169856 | 1.821536216 | 63.70334275 | 2.348308261 | 10.94391211 | 207.1543769 | 6.081090302 | 1.020684726 | 35.95255188 | 215.0195784 | 21.55761728 | 34.40404527 | 302.6542791 |
| E3-242-H | APOE3 | Male | 3 | 22.71977577 | 0 | 32.15226419 | 111.7261782 | 11.02004181 | 34.44725488 | 1.498975539 | 49.76809156 | 1.049103527 | 12.71075271 | 402.4022012 | 6.704982157 | 0.467277793 | 38.21245426 | 215.4167858 | 180.5185227 | 28.23210998 | 305.1620241 |
| E3-243-H | APOE3 | Male | 3 | 29.39605808 | 5.183691453 | 28.1859567 | 90.47812718 | 9.846665187 | 25.9900636 | 0.980243462 | 61.41362066 | 0.737491437 | 0.176646756 | 307.2547887 | 3.683084136 | 0 | 30.17203269 | 132.7469117 | 0 | 31.28020967 | 293.6090368 |
| E3-244-H | APOE3 | Male | 3 | 34.4240181 | 6.027872487 | 14.55770448 | 29.3976744 | 6.619749314 | 13.93291706 | 0.50983591 | 52.20989385 | 0.35716685 | 3.850918875 | 603.6429288 | 1.462601016 | 0 | 28.60168938 | 158.1066732 | 442.9248971 | 27.41492395 | 304.5766257 |
| E3-251-H | APOE3 | Male | 3 | 18.90959464 | 3.861853214 | 24.15351243 | 90.65793634 | 14.13422856 | 29.83721828 | 0.992477831 | 63.65744818 | 0.692762454 | 2.196171241 | 570.21181 | 2.507836055 | 0 | 21.50311272 | 122.2465837 | 318.4552386 | 27.67412569 | 301.2392854 |
| E3-252-H | APOE3 | Male | 3 | 15.17549072 | 0 | 25.72144955 | 95.1790814 | 10.65231214 | 50.332105 | 0.208064385 | 44.34111824 | 0.152618288 | 0 | 821.4541405 | 0.528597124 | 0 | 23.02853192 | 151.5605643 | 499.3744489 | 16.20935718 | 284.1499414 |
| E4-186-H | APOE4 | Male | 3 | 20.09241466 | 0 | 21.49051163 | 122.3539151 | 17.00140186 | 30.74260679 | 0.343518389 | 49.22823542 | 0 | 0 | 573.2452597 | 0.394138905 | 0 | 31.35149998 | 128.2513064 | 359.2805987 | 11.53586351 | 297.4539335 |
| E4-187-H | APOE4 | Male | 3 | 17.94403123 | 3.861853214 | 30.17203495 | 112.6271781 | 18.08553298 | 30.36051381 | 0.776662347 | 55.94870626 | 0.554596131 | 0 | 520.1370753 | 2.639162631 | 0 | 25.97441114 | 171.5548522 | 383.9202192 | 20.5203383 | 278.180083 |
| E4-188-H | APOE4 | Male | 3 | 20.48780869 | 0.985809465 | 21.28986474 | 125.8220728 | 14.48382064 | 38.58290372 | 1.915713812 | 45.67500543 | 0.349446397 | 6.697808444 | 358.4359438 | 7.15501173 | 0.817722136 | 34.71458203 | 151.9465919 | 125.1539582 | 17.91120827 | 303.5587255 |
| E4-189-H | APOE4 | Male | 3 | 21.64104062 | 8.056911029 | 23.23602871 | 120.5464673 | 16.11038816 | 34.34221131 | 2.028607768 | 48.84162835 | 1.747710863 | 9.258382971 | 354.7748474 | 8.683704984 | 2.421978456 | 32.95801851 | 135.4419775 | 245.7457754 | 32.0368555 | 301.2729025 |
| E4-198-H | APOE4 | Male | 3 | 19.19399622 | 0 | 12.7626507 | 101.124793 | 6.846429396 | 24.12001887 | 0.858994736 | 52.17598818 | 0.386377186 | 0 | 313.348619 | 3.085125496 | 0 | 20.78421886 | 123.4475645 | 0 | 29.0040518 | 296.9150545 |
| E4-199-H | APOE4 | Male | 3 | 18.52678724 | 0 | 23.94951972 | 137.8374578 | 14.85164491 | 32.8850612 | 0.397201503 | 60.71359086 | 0.422356499 | 0 | 372.2538676 | 2.023029522 | 0 | 27.77309767 | 175.4842498 | 0 | 22.39429161 | 287.4141262 |
| E4-200-H | APOE4 | Male | 3 | 16.51664287 | 2.192552978 | 15.20188757 | 94.93429503 | 13.19014677 | 24.44282637 | 0.951779249 | 48.51321646 | 0.218725697 | 0 | 254.2880528 | 0.372238822 | 0 | 21.93407865 | 168.9937066 | 0 | 27.88856122 | 300.5092356 |
| E4-210-H | APOE4 | Male | 3 | 21.58433579 | 2.638869052 | 8.585514924 | 148.4283204 | 16.26766389 | 31.31493527 | 1.16793604 | 67.28242953 | 0.697034501 | 7.600186516 | 91.86909867 | 8.653173365 | 4.336655451 | 50.82658064 | 243.7220199 | 0 | 33.97411675 | 283.3253829 |
| E4-3-H | APOE4 | Male | 18 | 27.07840984 | 8.054800859 | 32.94460239 | 124.2294547 | 19.57114324 | 35.0954752 | 3.179380049 | 71.23998518 | 2.509048961 | 21.93152246 | 404.8153176 | 9.313427528 | 6.843590044 | 62.54720135 | 188.9186013 | 138.3198018 | 31.17581155 | 301.6831147 |
| E4-4-H | APOE4 | Male | 18 | 20.09438702 | 3.401119568 | 19.9377437 | 128.7997779 | 16.42456826 | 38.13860035 | 0.728389716 | 47.63934503 | 1.126257454 | 0 | 253.9345837 | 4.068403919 | 0 | 43.05862074 | 174.8227244 | 0 | 26.13436702 | 294.924149 |
| E4-5-H | APOE4 | Male | 18 | 19.13068823 | 0 | 26.21350685 | 102.2078007 | 10.63503872 | 30.932893 | 0.00327854 | 51.58350487 | 0.109980631 | 0 | 313.2924674 | 1.395226761 | 0 | 32.92287265 | 140.7962429 | 0 | 13.25514626 | 279.9302696 |
| E4-6-H | APOE4 | Male | 18 | 16.96218831 | 3.900730241 | 29.7085217 | 110.9177898 | 9.303344817 | 20.10282975 | 0.686874448 | 51.65642825 | 0.5240893 | 1.033724481 | 226.6212041 | 2.305321689 | 0 | 39.97184108 | 101.6181185 | 0 | 14.31863745 | 293.4077882 |
| E4-29-H | APOE4 | Male | 18 | 20.41711927 | 0 | 24.69621502 | 157.3964451 | 17.28158578 | 56.1424849 | 0.237622062 | 47.33167688 | 1.16583411 | 0 | 445.7763892 | 2.268173536 | 2.271743003 | 115.5138835 | 223.6152302 | 126.1880688 | 15.22062329 | 282.7923071 |
| E4-30-H | APOE4 | Male | 18 | 21.30648205 | 2.192552978 | 34.97624695 | 102.2623005 | 20.84693849 | 34.6069356 | 3.134327756 | 48.41121763 | 1.801385162 | 14.9197364 | 362.2552563 | 4.888995976 | 3.403946299 | 110.534515 | 196.5830422 | 128.2900263 | 20.52098295 | 299.2301956 |
| E4-31-H | APOE4 | Male | 18 | 21.80119305 | 0.985809465 | 25.6161427 | 103.5406288 | 17.35122622 | 38.42788079 | 2.266037625 | 59.37987203 | 2.301510878 | 15.25407742 | 415.777993 | 6.736468621 | 2.011330088 | 56.51878027 | 204.5356726 | 0 | 20.55922335 | 300.1592629 |
| E4-32-H | APOE4 | Male | 18 | 15.59532359 | 2.423007423 | 21.61420482 | 112.6271781 | 8.775791209 | 36.05642109 | 0.877142791 | 59.8491117 | 1.399406781 | 0 | 225.988474 | 4.283199167 | 0 | 46.58397639 | 169.9308626 | 57.62529513 | 25.33775788 | 293.6090368 |
| E3-20-H | APOE3 | Male | 18 | 21.95994157 | 5.229556537 | 33.33125954 | 184.3815988 | 10.09919143 | 45.06752058 | 1.870350396 | 49.74368503 | 1.966208903 | 8.769439745 | 318.3209515 | 6.298131417 | 6.530306786 | 47.27408677 | 174.7818034 | 257.1774509 | 14.98940222 | 288.9792368 |
| E3-44-H | APOE3 | Male | 18 | 20.55125331 | 8.724781876 | 18.62650496 | 115.4655857 | 11.96537315 | 24.26087165 | 1.57707789 | 49.39856773 | 2.497056359 | 13.81409456 | 427.7680402 | 7.03677666 | 0 | 49.17215864 | 177.4818925 | 112.5923314 | 39.66292497 | 308.6088559 |
| E3-60-H | APOE3 | Male | 18 | 22.36827029 | 5.043206526 | 28.14735773 | 105.6063101 | 15.65588487 | 33.92050502 | 0.947375742 | 68.65809609 | 2.294419013 | 10.90488971 | 299.9400022 | 6.849867372 | 0 | 51.93727033 | 172.9498216 | 0 | 38.96394871 | 290.9772778 |
| E3-61-H | APOE3 | Male | 18 | 19.03010377 | 1.669300237 | 31.16607874 | 101.4820193 | 15.04352052 | 34.42848805 | 1.367181592 | 57.39732786 | 0.433382706 | 0.176646756 | 225.1314865 | 5.255148738 | 0 | 56.57555691 | 171.2908264 | 0 | 32.71161541 | 294.7593824 |
| E3-62-H | APOE3 | Male | 18 | 18.18209548 | 0.985809465 | 22.27110895 | 77.74262151 | 14.97400352 | 31.62608296 | 0.232238365 | 56.32542966 | 1.034737748 | 4.255253822 | 133.9691028 | 6.286978682 | 0 | 50.64518724 | 173.8754293 | 0 | 28.05607292 | 306.8773076 |
| E3-92-H | APOE3 | Male | 18 | 22.06619115 | 3.900730241 | 20.0484516 | 181.4205543 | 8.704472657 | 49.49556271 | 1.653164966 | 50.82380143 | 1.047871047 | 8.100637903 | 241.3079497 | 6.010701343 | 10.75125535 | 76.48348293 | 232.1270019 | 11.58086998 | 21.31270109 | 308.2510263 |
| E3-93-H | APOE3 | Male | 18 | 22.2237619 | 8.770411132 | 17.74823282 | 129.5595537 | 11.2966911 | 40.08650723 | 2.132806545 | 52.54731147 | 3.47657017 | 17.20611699 | 413.5075656 | 7.215257428 | 4.505518401 | 63.72990284 | 244.8868672 | 438.6428329 | 31.58752179 | 308.2996728 |
| E3-101-H | APOE3 | Male | 18 | 22.98356706 | 0.458407707 | 14.82914837 | 117.56393 | 6.320694533 | 48.68878271 | 0.790647479 | 55.06078913 | 0.513748166 | 1.932659172 | 464.6505573 | 4.752515051 | 16.68042977 | 103.1375386 | 188.9146216 | 0 | 22.59957564 | 282.7263963 |
